# An ASXL3–thyroid hormone axis in parvalbumin interneurons controls autism-like behaviors

**DOI:** 10.64898/2026.05.22.727319

**Authors:** Chaodong Ding, Yiting Yuan, Yixin Hu, Yuefang Zhang, Shihao Wu, Ailian Du, Zilong Qiu

## Abstract

Pathogenic variants in *ASXL3* underlie autism spectrum disorder (ASD), but how they perturb neural circuits and whether defects are reversible remain unclear. Here we show that *Asxl3* haploinsufficiency in mice reduces cortical thickness and upper-layer projection neurons while increasing parvalbumin (PV) interneuron density and producing ASD-like behavioral abnormalities. Mechanistically, *Asxl3* loss derepresses the thyroid hormone (TH)–inactivating enzyme DIO3 via altered histone H2A monoubiquitination, depleting brain TH. Conditional deletion of the TH receptor *Thra* in inhibitory neuron progenitors phenocopies the PV interneuron expansion, linking impaired TH signaling to PV circuit remodeling. Neonatal, but not adolescent, TH supplementation restores PV interneuron numbers and rescues behavior in *Asxl3^+/−^* mice, defining a critical early window for intervention. An intein-based AAV system that reconstitutes full-length ASXL3 normalizes cortical architecture and behavior in *Asxl3^+/−^* mice and drives efficient ASXL3 expression in non-human primate brain, establishing an ASXL3–TH–PV interneuron axis as a targetable pathway in ASD.

## Introduction

Autism spectrum disorder (ASD) is a highly heterogeneous neurodevelopmental condition characterized by impairments in social communication, restricted interests, and repetitive behaviors, and is frequently accompanied by intellectual disability, epilepsy, and anxiety^1^. The disorder typically manifests in early childhood and is more prevalent in males^2^. Genetic studies indicate that *de novo* and inherited variants are major contributors to ASD risk^3^, and large-scale sequencing efforts have identified hundreds of ASD-associated genes involved in synaptic function, chromatin regulation, and neurodevelopmental programs^4–8^. However, for many risk genes, the relevant neural cell types, developmental windows, and molecular pathways that link mutation to circuit dysfunction and behavior remain poorly defined.

Among candidate ASD risk genes, *ASXL3* has emerged as a recurrently mutated locus in pediatric cohorts^9–12^. *ASXL3* encodes a core component of the polycomb deubiquitinase complex that removes monoubiquitin from histone H2A at lysine 119, thereby modulating chromatin accessibility and transcriptional programs^13^. ASXL3 is highly expressed in the developing brain and is required for normal embryonic development and neurogenesis, consistent with the severe neurodevelopmental phenotypes observed in individuals carrying *ASXL3* loss-of-function variants^14, 15^. These observations highlight *ASXL3* as a strong ASD risk gene, but how *ASXL3* haploinsufficiency remodels neural circuits and whether the resulting defects are reversible remain unknown.

Converging evidence points to PV-expressing interneurons as a key cellular substrate in ASD^16^. These fast-spiking GABAergic interneurons are crucial for maintaining excitation–inhibition balance and generating network oscillations that support social behavior, cognitive flexibility, and memory^17–19^. Altered number, distribution, or function of PV interneurons has been reported in postmortem ASD cortex and in multiple genetic mouse models^20–22^, suggesting that PV interneuron dysfunction is a common downstream node for diverse ASD-associated perturbations. Yet, the upstream genetic and molecular mechanisms that govern PV interneuron development in the context of ASD risk genes, including epigenetic regulators such as ASXL3, remain largely undefined.

TH signaling represents another axis implicated in ASD pathogenesis. Epidemiological studies link maternal hypothyroidism and other TH metabolism abnormalities to increased ASD risk in offspring^23–25^, and experimental work indicates that TH directly regulates neural progenitor proliferation and neuronal differentiation^26, 27^. TH insufficiency during early postnatal development disrupts the maturation of PV interneurons in the cortex and hippocampus^28^, and TH acts primarily through the nuclear receptors THRA and THRB, with GABAergic neurons, including PV interneurons, as major targets^29, 30^. These observations suggest that genetic disruptions and hormonal imbalance may converge on TH-dependent developmental programs in inhibitory circuits. However, whether ASD-associated mutations in *ASXL3* perturb TH signaling, how this affects PV interneuron development, and whether such defects can be corrected within a defined developmental window have not been systematically explored.

Here, we identify an ASXL3–TH–PV interneuron axis that links epigenetic dysregulation to circuit remodeling and ASD-like behaviors. Using *Asxl3* haploinsufficient (*Asxl3*^+/-^) mice, we show that reduced *Asxl3* expression leads to cortical thinning, loss of upper-layer projection neurons, expansion of PV interneuron populations in retrosplenial cortex (RSC) and hippocampus, and robust ASD-like behavioral abnormalities. At the molecular level, *Asxl3* loss derepresses the expression of the TH-inactivating enzyme gene *Dio3* via regulation of histone H2A monoubiquitination at its promoter, resulting in reduced TH levels in serum and brain. Consistent with a causal role for impaired TH signaling in inhibitory lineages, conditional deletion of the TH receptor *Thra* in Nkx2.1-expressing inhibitory neuron progenitors increases PV interneuron density, phenocopying the *Asxl3*^+/-^ phenotype. Neonatal, but not adolescent, T₃ supplementation restores TH levels, normalizes PV interneuron numbers, and rescues ASD-like behaviors in *Asxl3*^+/-^ mice, revealing a critical early-life window for intervention. Finally, using an intein-based AAV strategy to reconstitute full-length ASXL3, we demonstrate that restoring ASXL3 expression in neonatal *Asxl3*⁺/⁻ mice reverses cortical and hippocampal neurodevelopmental defects and normalizes behavior, and we establish efficient full-length ASXL3 expression in the brains of non-human primates. Together, these findings define an epigenetic–hormonal–circuit axis in ASD pathogenesis and suggest that neonatal TH supplementation and *ASXL3*-targeted gene restoration are promising therapeutic strategies for *ASXL3*-related and potentially broader forms of ASD.

## Results

### *Asxl3* haploinsufficiency reduces cortical thickness and selectively alters neuronal populations in mice

In our previous study, we identified *de novo* nonsense mutations in *ASXL3* in two Chinese children with ASD (Fig. 1A) ^12^. To begin probing *ASXL3* function in the brain, we first examined its spatial and temporal expression patterns. Analysis of public RNA-seq datasets showed that *ASXL3* is highly expressed in brain, ovary, and testis, with particularly enriched expression in fetal compared with postnatal brain tissue (Fig. S1A, S1B). Within the human brain, *ASXL3* is broadly expressed across regions and is especially abundant in the cortex (Fig. S1C). Single-cell transcriptomic data further revealed *ASXL3* expression in multiple classes of excitatory and inhibitory neurons in the human cortex (Fig. S1D, S1E).

**Figure 1.**
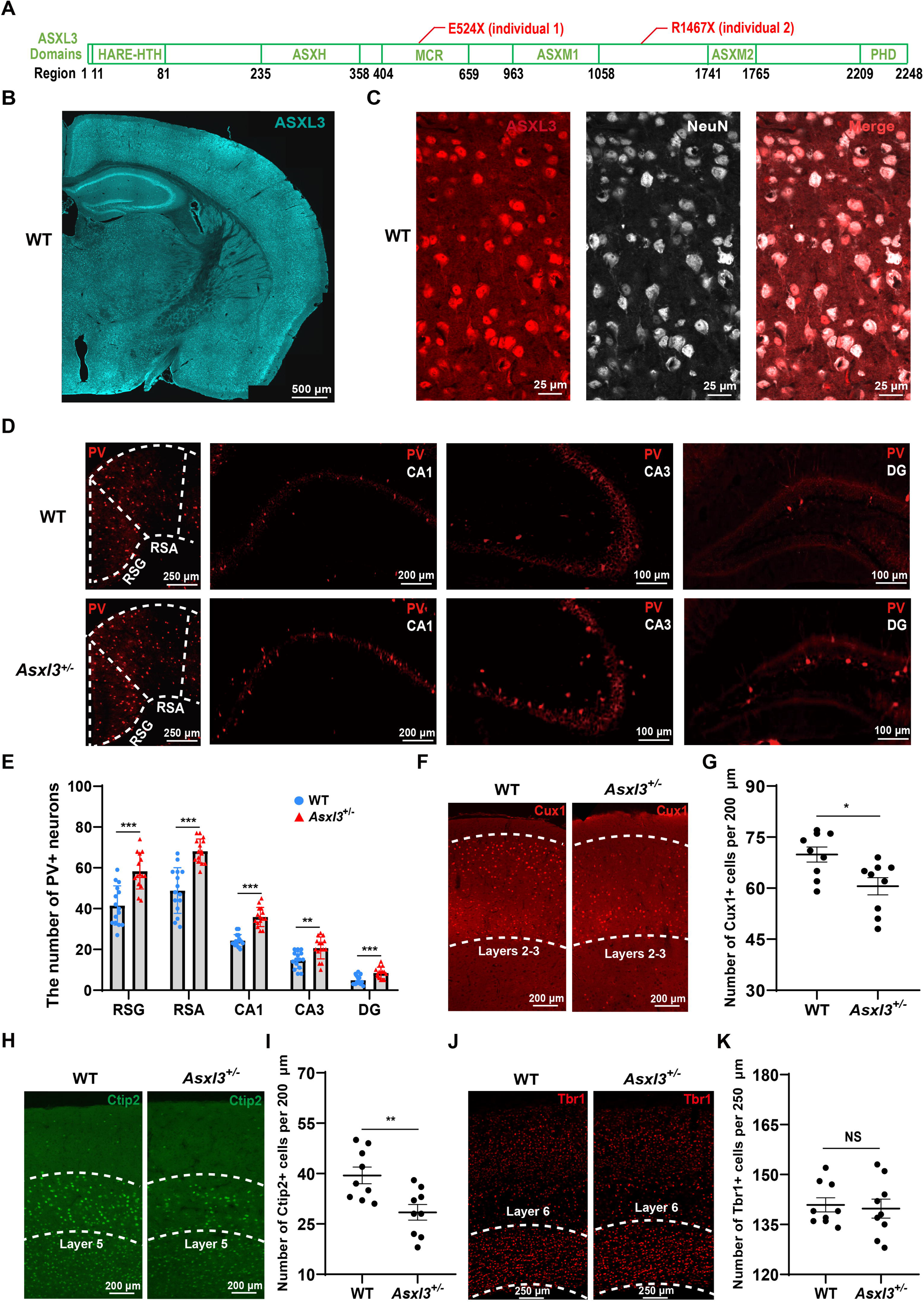
Heterozygous knockout of *Asxl3* increases PV+ neurons and decreases upper-layer cortical neurons. (A) Identification of two *de novo* nonsense mutations in *ASXL3* from Chinese children with ASD. (B) Immunofluorescence staining of a mouse brain section using an ASXL3-specific antibody. (C) Double immunofluorescence staining of the mouse cortex with ASXL3 and NeuN antibodies. (D) and (E) Immunostaining for PV-positive neurons in the RSC and hippocampus, accompanied by quantification of their counts (n = 15 slices from 5 mice). The RSC encompasses the RSG and RSA regions. (F) and (G) Immunostaining for Cux1-positive neurons in the cortex, with quantification of their counts (n = 9 slices from 3 mice). Cux1 is a marker for neurons in the second and third cortical layers. (H) and (I) Immunostaining for Ctip2-positive neurons in the cortex, with quantification of their counts (n = 9 slices from 3 mice). Ctip2 is a marker for neurons in the fifth cortical layer. (J) and (K) Immunostaining for Tbr1-positive neurons in the cortex, with quantification of their counts (n = 9 slices from 3 mice). Tbr1 is a marker for neurons in the sixth cortical layer. Data are depicted as mean ± SD, with statistical significance determined by a two-tailed student’s t-test, \**P <* 0.05, \*\**P <* 0.01, \*\*\**P <* 0.001, NS (not significant).

Consistent with the human data, *Asxl3* is widely expressed in the mouse brain and is largely neuronal (Fig. 1B, 1C). Mouse ENCODE transcriptomic data (BioProject PRJNA66167) confirmed that *Asxl3* expression is higher in brain than in other tissues and is enriched in embryonic relative to adult brain (Fig. S1F). Single-cell RNA-seq from the Allen Brain Map demonstrated *Asxl3* expression across diverse inhibitory neurons, excitatory neurons, and glial populations (Fig. S1G), supporting a broad role for *Asxl3* in neurodevelopment.

To model *ASXL3* haploinsufficiency *in vivo*, we generated heterozygous *Asxl3* knockout (*Asxl3*^+/-^) mice using CRISPR/Cas9 to delete exons 5–14 of *Asxl3* (Fig. S2A, S2B). PCR-based genotyping identified *Asxl3*^+/-^ animals (Fig. S2B, S2C), which displayed an approximately 50% reduction of *Asxl3* mRNA and protein in the brain compared with wild-type (WT) littermates (Fig. S2D–S2F). *Asxl3*^+/-^ mice exhibited reduced body weight, brain size, and brain weight (Fig. S3A–S3D), and Nissl staining revealed a significant decrease in cortical thickness (Fig. S3E, S3F), indicating that *Asxl3* haploinsufficiency impairs brain growth and cortical development.

We next asked which neuronal populations are preferentially affected by *Asxl3* loss. Immunofluorescence staining revealed a marked increase in the number of PV-positive interneurons in the RSC (including RSG and RSA) and hippocampus of *Asxl3*^+/-^ mice relative to WT controls (Fig. 1D, 1E). By contrast, the numbers of somatostatin (SST)-positive and vasoactive intestinal polypeptide (VIP)-positive interneurons were unchanged between genotypes (Fig. S4A–S4H). Glial cell composition also appeared preserved: we detected no significant differences in microglia or oligodendrocyte numbers in the cortex or hippocampus of *Asxl3*^+/-^ mice (Fig. S4I–S4P). These findings indicate that *Asxl3* haploinsufficiency selectively alters PV interneuron abundance without broadly affecting other interneuron subtypes or glial populations.

Given the reduction in cortical thickness, we further examined whether specific cortical layers and projection neuron populations were affected. Staining for Cux1, a marker of layer 2/3 neurons, revealed a significant reduction in Cux1-positive neurons in the superficial layers of *Asxl3*^+/-^ cortex (Fig. 1F, 1G). We also observed a decrease in Ctip2-positive layer 5 neurons in *Asxl3*^+/-^ mice compared with WT (Fig. 1H, 1I), whereas the number of Tbr1-positive layer 6 neurons was unchanged (Fig. 1J, 1K). Together, these results show that *Asxl3* haploinsufficiency leads to cortical thinning accompanied by loss of upper-layer and layer 5 projection neurons and a concomitant increase in PV-positive interneuron density in specific cortical and hippocampal regions.

### *Asxl3* deficiency induces ASD-like phenotypes in mice

We first assessed social behavior in male *Asxl3*^+/-^ mice using the three-chamber test. During the habituation phase, WT and *Asxl3*^+/-^ mice showed no side preference when both chambers were empty (Fig. S5A, S5B). In the sociability phase, both genotypes spent more time interacting with a stranger mouse than with an empty cage (Fig. 2A, 2B), indicating preserved basic sociability. In contrast, in the social novelty phase, WT mice preferentially interacted with a novel stranger over a familiar mouse, whereas *Asxl3*^+/-^ mice failed to show this preference (Fig. 2C, 2D), revealing a specific deficit in social novelty recognition.

**Figure 2.**
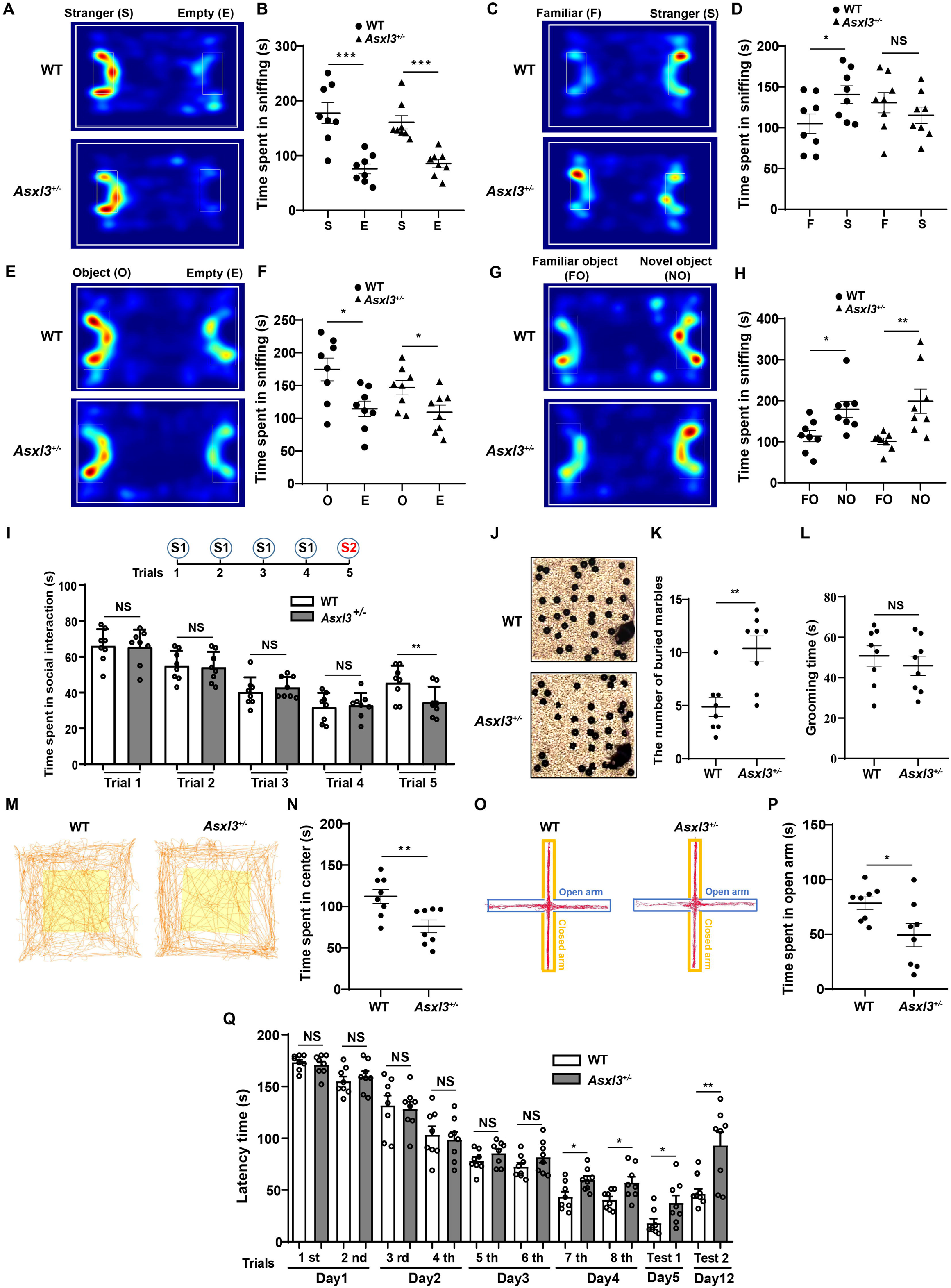
The *Asxl3*^+/-^ mice exhibited ASD-like behavioral phenotypes. (A) and (B) Tracing heatmaps and quantification of social interaction time in the sociability test (C) and (D) Tracing heatmaps and quantification of social interaction time in the social novelty test. (E) and (F) Tracing heatmaps and time spent exploring objects in the object cognition test. (G) and (H) Tracing heatmaps and time spent exploring objects in the novel object recognition test. (I) Social interaction time between the tested mice and the introduced mice in the social intruder test. (J) and (K) Representative images and quantification of buried marbles in the marble burying test. (L) Quantification of self-grooming time for WT and *Asxl3*^+/-^ mice. (M) and (N) Representative traces and time spent in the central area of the open field test. (O) and (P) Representative traces and time spent in the open arms of the elevated plus maze test. (Q) Statistics on the time it took mice to find the escape hole in the Barnes maze test. All data are presented as mean ± SD, with n = 8 for both WT and *Asxl3*^+/-^ mice. Statistical significance was determined using a two-tailed student’s t-test, where \**P <* 0.05, \*\**P <* 0.01, \*\*\**P <* 0.001, and NS indicates not significant.

To test whether this impairment extended to non-social recognition, we performed the novel object recognition task. As in the three-chamber task, neither genotype preferred either side during the initial habituation with empty cages (Fig. S5C, S5D). When objects were introduced, both WT and *Asxl3*^+/-^ mice spent more time exploring objects than empty cages (Fig. 2E, 2F). Upon the introduction of a novel object, both groups exhibited a preference for the new object over the familiar one (Fig. 2G, 2H), indicating intact object recognition memory in male *Asxl3*^+/-^ mice and dissociation between social and non-social novelty processing.

We next used the social intruder test to further probe social novelty. Across the first four trials with the same intruder, WT and *Asxl3*^+/-^ males showed comparable interaction times (Fig. 2I). However, when a new intruder was introduced in the fifth trial, *Asxl3*^+/-^ mice spent significantly less time interacting compared with WT littermates (Fig. 2I), corroborating a selective deficit in recognizing and engaging with novel social partners.

We then examined core ASD-related behavioral domains beyond sociability. In the marble burying test, *Asxl3*^+/-^ males buried more marbles than WT controls (Fig. 2J, 2K), consistent with increased repetitive and stereotyped behavior. Self-grooming duration did not differ between genotypes (Fig. 2L), suggesting that repetitive phenotypes may be assay-dependent. Anxiety-like behavior was elevated in *Asxl3*^+/-^mice: in the open field, they spent less time in the central area (Fig. 2M, 2N), and in the elevated plus maze they spent less time in the open arms (Fig. 2O, 2P). Finally, in the Barnes maze test, *Asxl3*^+/-^ males required more time to locate the escape hole than WT mice (Fig. 2Q), indicating impaired spatial learning and memory.

To determine whether these phenotypes are sex-independent, we repeated the behavioral battery in female *Asxl3*^+/-^ mice. Similar deficits in social and object recognition were observed in the three-chamber and novel object recognition tests, respectively (Fig. S5E-S5H), suggesting broader recognition memory impairment in females. In the social intruder test, female *Asxl3*^+/-^ mice also demonstrated deficits in recognizing new social partners (Fig. S5I).

As in males, female *Asxl3*^+/-^ mice displayed increased marble burying, heightened anxiety-like behavior in the open field and elevated plus maze, and longer escape latencies in the Barnes maze test (Fig. S5J–S5P). Together, these results demonstrate that *Asxl3* haploinsufficiency induces robust ASD-like behavioral abnormalities—including social novelty deficits, enhanced repetitive behavior, increased anxiety, and impaired spatial learning—in both male and female mice.

### *Asxl3* downregulation upregulates the thyroid hormone–inactivating enzyme DIO3 and reduces brain TH levels

To identify downstream targets through which *Asxl3* haploinsufficiency might influence neurodevelopment, we performed RNA sequencing on mixed cortical and hippocampal tissue from *Asxl3*^+/-^ and WT mice. Differential expression analysis revealed 837 genes whose expression was significantly altered in *Asxl3*^+/-^ brains, with 435 upregulated and 402 downregulated transcripts (Fig. 3A). Gene Ontology enrichment analysis indicated that these genes are involved in key neurodevelopmental processes, including neurogenesis, neuronal differentiation, and axonal projection (Fig. 3B). From the differentially expressed genes, we selected ten candidates with robust changes and neurodevelopmental relevance for validation. RT-qPCR confirmed these alterations and highlighted *Dio3* as the most strongly upregulated gene in both cortex and hippocampus of *Asxl3*^⁺/⁻^ mice (Fig. 3C, 3D). Consistently, DIO3 protein levels were increased in the cortex of *Asxl3*^⁺/⁻^ mice compared with WT littermates (Fig. 3E, 3F), suggesting that *Asxl3* downregulation enhances expression of the TH–inactivating enzyme DIO3.

**Figure 3.**
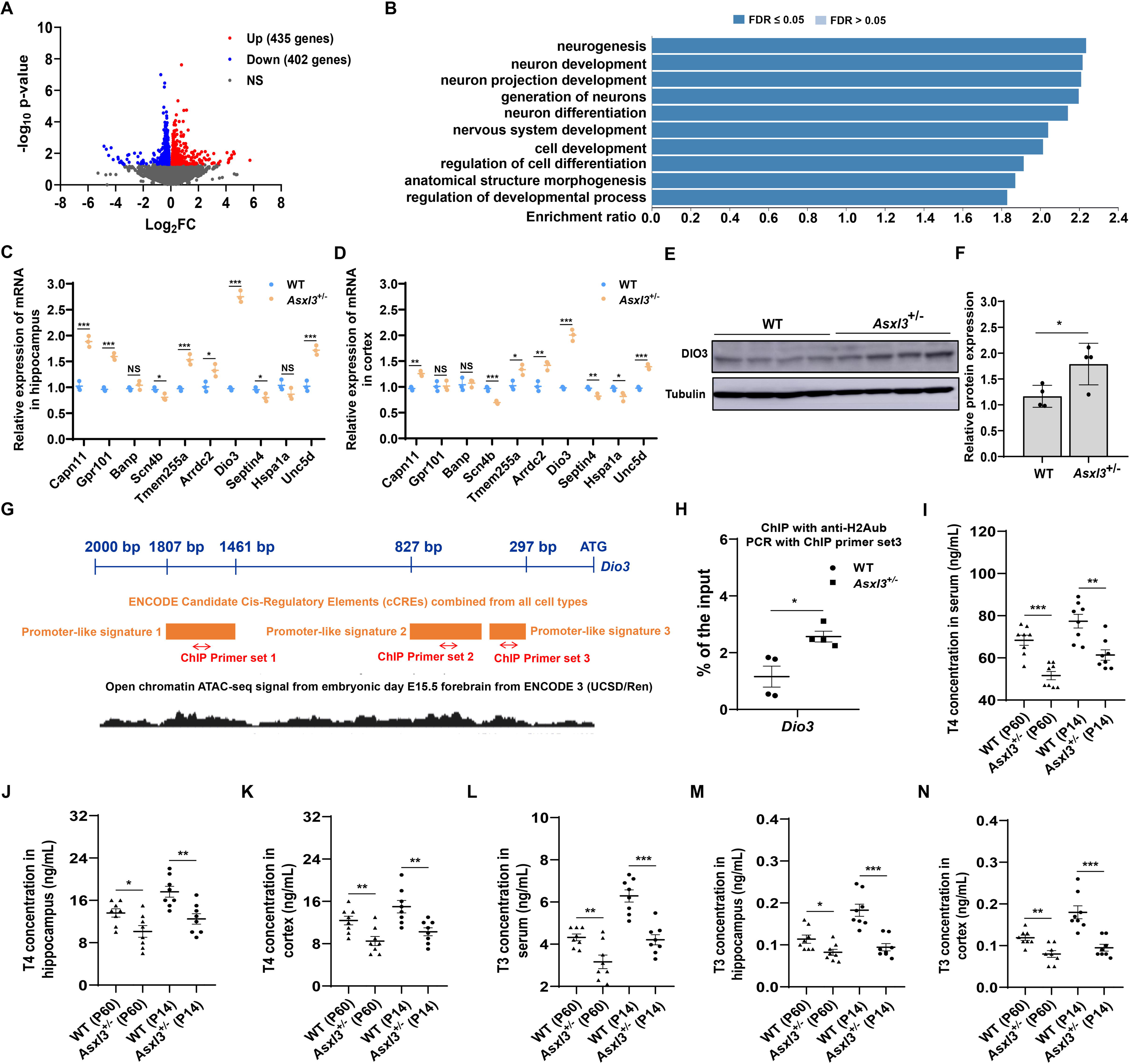
*Asxl3* regulates the expression of thyroxine metabolism-related gene Dio3 **(A)** The volcano plot illustrates the RNA sequencing outcomes, highlighting differential gene expression (n = 3 mice per group). (B) Functional enrichment analysis delineates the biological processes associated with the 837 significantly expressed genes. (C) and (D) RT-qPCR was utilized to ascertain the expression levels of the *Asxl3* target genes in the hippocampal and cortical regions (n = 3 mice per group). (E) and (F) The protein expression levels of *Dio3* in both WT and *Asxl3*^+/-^ mice were measured and quantified (n = 4 mice per group). (G) Schematic representation of the promoter regions upstream of the *Dio3* alongside the ATAC-seq signals. (H) ChIP assay was conducted to evaluate the binding of H2Aub at the *Dio3* promoter in WT and *Asxl3*^+/-^ mice (n = 4 mice per group). (I) - (K) ELISA was employed to measure T4 levels in serum, hippocampus, and cortex of WT and *Asxl3*^+/-^ mice aged 2 weeks and 2 months (n = 8 mice per group). (L) - (N) ELISA was employed to measure T3 levels in serum, hippocampus, and cortex of WT and *Asxl3*^+/-^ mice aged 2 weeks and 2 months (n = 8 mice per group). Data are presented as mean ± SD. Statistical significance was assessed using a two-tailed student’s t-test, with \**P <* 0.05, \*\**P <* 0.01, \*\*\**P <* 0.001 indicating significance, and NS denoting no significance.

Given the epigenetic role of ASXL3, we next examined how *Asxl3* regulates *Dio3* at the chromatin level. Inspection of public ATAC-seq profiles using the UCSC Genome Browser identified three putative regulatory regions upstream of *Dio3* marked by open chromatin, indicative of promoter or enhancer activity (Fig. 3G). We designed ChIP-qPCR primers against these regions and assessed H2A monoubiquitination (H2Aub) using an H2Aub-specific antibody. Among these three regions, only the third amplicon displayed a ChIP signal, and a significant enrichment of H2Aub was observed at the *Dio3* promoter in *Asxl3*^+/-^ mice (Fig. 3H).

Because DIO3 catalyzes TH inactivation, we next asked whether *Dio3* upregulation in *Asxl3*^+/-^ mice impacts systemic and brain TH levels. ELISA measurements in 2-week-old and 2-month-old mice showed significantly reduced thyroxine (T4) and triiodothyronine (T3) levels in serum, cortex, and hippocampus of *Asxl3*^+/-^ mice compared with WT controls (Fig. 3I–3N). Together, these data identify *Dio3* as a key *Asxl3*-regulated target and demonstrate that *Asxl3* downregulation leads to *Dio3* upregulation and a concomitant reduction of TH levels in the developing brain.

### *Thra* deletion in inhibitory neuron progenitors increases PV interneuron density

Because *Asxl3* haploinsufficiency reduces TH levels and increases PV interneuron density, we next asked whether dampening TH signaling specifically in inhibitory neuron progenitors is sufficient to drive PV interneuron expansion. To this end, we conditionally deleted the TH receptor *Thra* in Nkx2.1-expressing inhibitory neuron progenitors by crossing Nkx2.1-CreER mice with *Thra*^fl/+^ mice to generate Nkx2.1-*Thra*^fl/+^ animals. Newborn Nkx2.1-*Thra*^fl/+^ pups received intraperitoneal tamoxifen injections for three consecutive days to induce Cre activity in Nkx2.1-positive progenitors (Fig. 4A).

**Figure 4.**
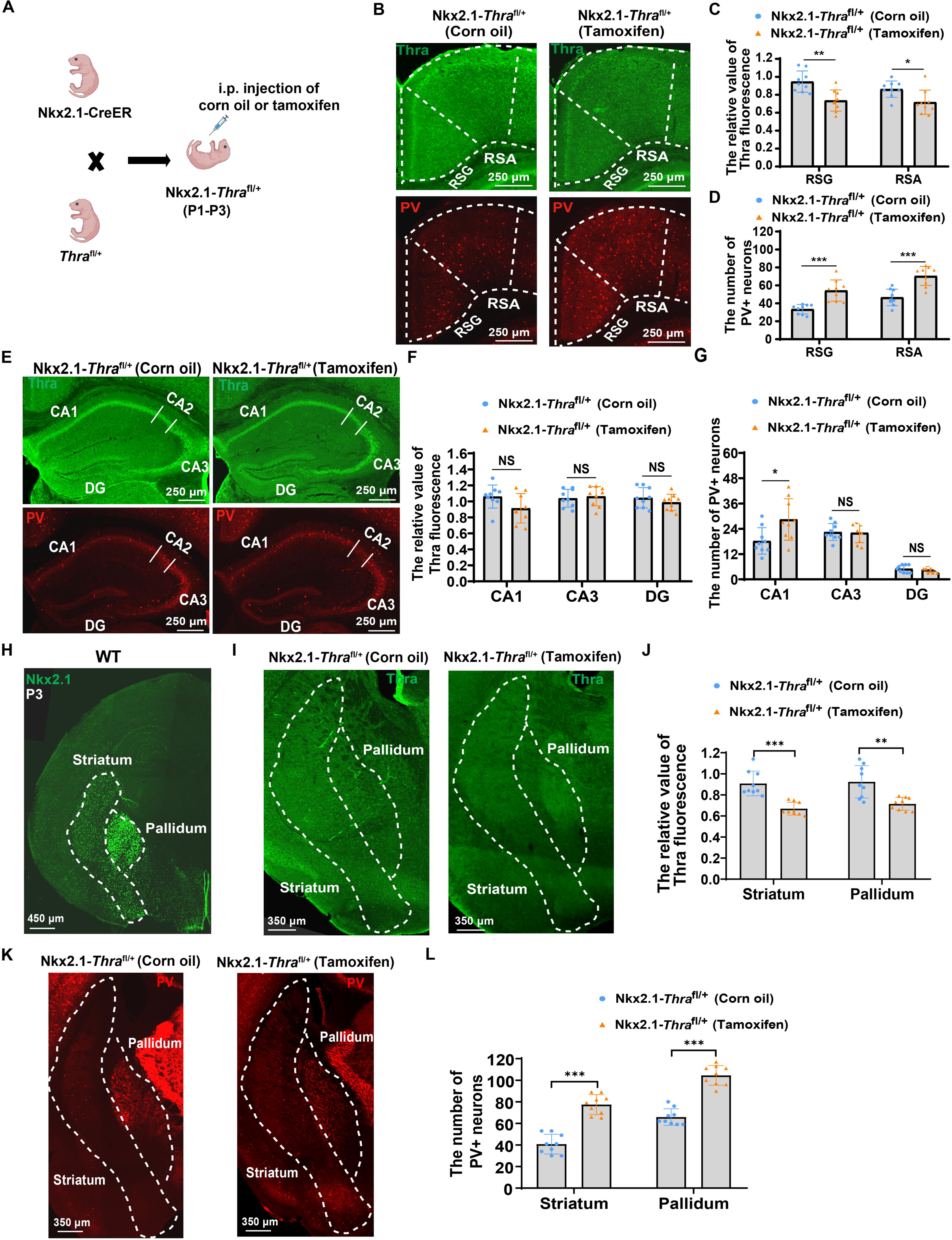
Effects of specific deletion of *Thra* in inhibitory neuron progenitors on the development of pv-positive neurons. (A) Tamoxifen was administered to Nkx2.1-*Thra*^fl/+^ mice to induce *Thra*-specific knockout in inhibitory neuron progenitors. Tamoxifen was administered at a dose of 75 ng/g every 24 hours for 3 consecutive days, with corn oil injection serving as the control. (B) Immunofluorescent staining for Thra and PV in the RSC of Nkx2.1-*Thra*^fl/+^ mice after corn oil or tamoxifen injection. (C) Quantification of Thra fluorescence intensity in the RSC of Nkx2.1-*Thra*^fl/+^ mice. (D) Quantification of PV-positive neuron number in the RSC of Nkx2.1-*Thra*^fl/+^ mice. (E) Immunofluorescent staining for Thra and PV in the hippocampus of Nkx2.1-*Thra*^fl/+^ mice after corn oil or tamoxifen injection. (F) Quantification of Thra fluorescence intensity in the hippocampus of Nkx2.1-*Thra*^fl/+^ mice. (G) Quantification of PV-positive neuron number in the hippocampus of Nkx2.1-*Thra*^fl/+^ mice. (H) Immunofluorescent staining for Nkx2.1 in brain sections from WT mice at postnatal day 3. (I) Immunofluorescent staining for Thra in the striatum and globus pallidus of Nkx2.1-*Thra*^fl/+^ mice after corn oil or tamoxifen injection. (J) Quantification of Thra fluorescence intensity in the striatum and globus pallidus of Nkx2.1-*Thra*^fl/+^ mice. (K) Immunofluorescent staining for PV in the striatum and globus pallidus of Nkx2.1-*Thra*^fl/+^ mice after corn oil or tamoxifen injection. (L) Quantification of PV-positive neuron number in the striatum and globus pallidus of Nkx2.1-*Thra*^fl/+^ mice. Data are expressed as mean ± SD, n = 9 slices from 3 mice. Statistical analysis was performed using two-tailed Student’s t-test. \**P* < 0.05, \*\**P* < 0.01, \*\*\**P* < 0.001, NS (not significant).

At one month of age, we first examined *Thra* expression in the RSC. Compared with corn oil–injected controls, tamoxifen-treated Nkx2.1-*Thra*^fl/+^ mice showed a marked reduction in *Thra* signal in the RSC (Fig. 4B, 4C), accompanied by a significant increase in PV-positive interneuron number in this region (Fig. 4B, 4D). In the hippocampus, *Thra* signal was detectable in both groups and did not differ significantly at the population level (Fig. 4E, 4F). Nevertheless, PV-positive interneuron density in the CA1 region was significantly increased in tamoxifen-induced Nkx2.1-*Thra*^fl/+^ mice (Fig. 4E, 4G), indicating that reduced THRA function in Nkx2.1-lineage cells is sufficient to enhance PV interneuron numbers even in regions where bulk *Thra* levels appear largely preserved.

Because *Nkx2.1* is abundantly expressed in the striatum and globus pallidus of newborn mice (Fig. 4H), we next examined these nuclei. In tamoxifen-induced Nkx2.1-*Thra*^fl/+^ mice, *Thra* expression was significantly decreased in both striatum and globus pallidus compared with controls (Fig. 4I, 4J), and this reduction was accompanied by a robust increase in PV-positive interneuron number in both regions (Fig. 4K, 4L). Together, these data demonstrate that decreasing THRA-mediated TH signaling in Nkx2.1-lineage inhibitory neuron progenitors increases PV interneuron density across multiple brain regions, functionally linking TH signaling to PV interneuron development.

### Neonatal, but not adolescent, T3 supplementation rescues ASD-like phenotypes in *Asxl3^⁺/⁻^* mice

Given that *Asxl3* haploinsufficiency reduces brain T₃ levels and alters PV interneuron development, we next asked whether restoring T₃ during early life could rescue the structural and behavioral phenotypes in *Asxl3*^+/-^ mice. Neonatal *Asxl3*^+/-^ pups received intraperitoneal T_3_ injections every other day for 21 days (Fig. 5A). ELISA confirmed that this regimen significantly increased T_3_ levels in serum, cortex, and hippocampus (Fig. 5B). Strikingly, neonatal T_3_ supplementation normalized PV-positive interneuron density in the RSC and hippocampus (Fig. 5C–5F), indicating correction of the inhibitory circuit abnormalities.

**Figure 5.**
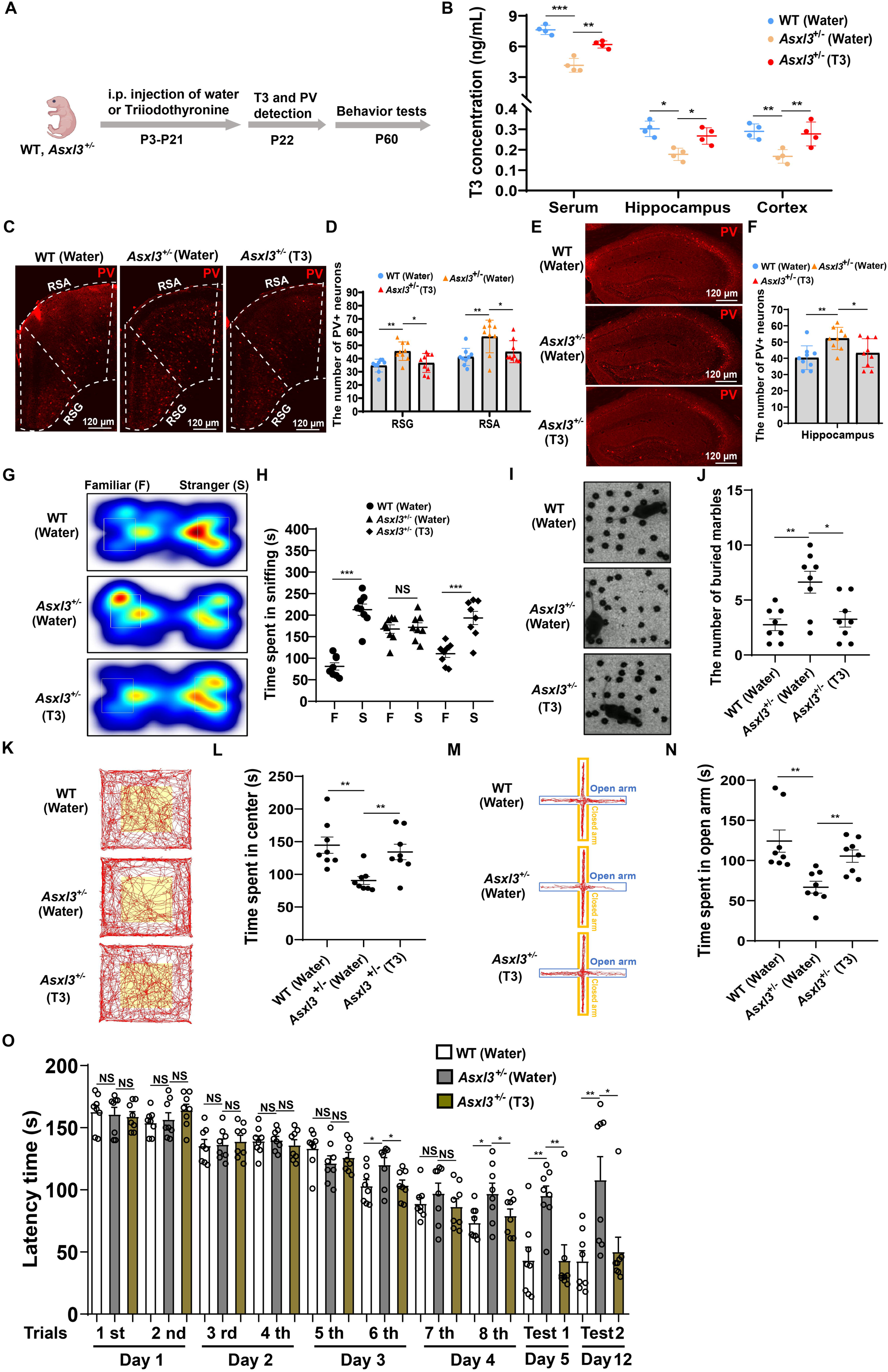
T3 supplementation in neonatal *Asxl3*^+/-^ mice rescued ASD-like phenotypes. (A) The workflow of T3 injection and subsequent phenotypic testing in *Asxl3*^+/-^ mice. T3 was administered to mice at a dose of 200 ng/g every other day from postnatal day 3 to postnatal day 21 (P3–P21), with normal saline injection serving as the control. (B) Expression of T3 in the serum and brain of *Asxl3*^+/-^ mice after T3 injection as determined by ELISA (n = 4 mice per group). (C) - (F) The examination and quantification of the expression of PV within the RSC and hippocampus following T3 supplementation in neonatal mice (n = 9 slices from 3 mice). (G) and (H) Representative heat maps of social novelty test results and statistics of social interaction time (n = 8 mice per group). (I) and (J) Graphical results of the marble burying test and quantitative analysis of the number of buried marbles (n = 8 mice per group). (K) and (L) Representative traces of the open field test results and statistics of the time mice spent exploring the central area (n = 8 mice per group). (M) and (N) Representative traces of the elevated plus maze test results and statistics of the time mice spent exploring the open arms (n = 8 mice per group). (O) Statistics of the time taken by mice to find the escape hole in the Barnes maze test (n = 8 mice per group). Data are presented as mean ± SD. Statistical significance was determined using a two-tailed student’s t-test for Fig. 5H and one-way ANOVA for Figs. 5B, 5D, 5F, 5J, 5L, 5N, 5O. \**P <* 0.05, \*\**P <* 0.01, \*\*\**P <* 0.001, NS (not significant).

We then examined whether these circuit-level improvements translated into behavioral rescue. In the three-chamber social novelty test, T₃-treated *Asxl3*^+/-^ mice spent more time interacting with a novel stranger mouse than with a familiar one (Fig. 5G, 5H), reflecting restored social novelty recognition. In the marble burying assay, T_3_ supplementation significantly reduced the number of buried marbles in *Asxl3*^+/-^mice (Fig. 5I, 5J), suggesting amelioration of repetitive, stereotyped behavior. Anxiety-like behavior was also improved: T₃-treated *Asxl3*^+/-^ mice spent more time in the center of the open field (Fig. 5K, 5L) and in the open arms of the elevated plus maze (Fig. 5M, 5N). Finally, in the Barnes maze, T₃-treated *Asxl3*^+/-^ mice showed reduced latency to locate the escape hole (Fig. 5O), indicating enhanced spatial learning and memory. Together, these data demonstrate that neonatal T₃ supplementation is sufficient to normalize PV interneuron development and broadly rescue ASD-like behavioral abnormalities in *Asxl3*^+/-^ mice.

To test whether this rescue is restricted to an early developmental window, we next supplemented T_3_ in adolescent *Asxl3*^+/-^ mice. T₃ was delivered in the drinking water from adolescence until 2 months of age (Fig. S6A), which significantly increased T_3_ levels in serum, hippocampus, and cortex (Fig. S6B). Despite effective hormonal supplementation, adolescent T_3_ treatment failed to correct the behavioral deficits: in the three-chamber social novelty test, *Asxl3*^+/-^ mice still did not show a preference for the novel over the familiar mouse (Fig. S6C, S6D), and in the marble burying test, T_3_ did not reduce the number of buried marbles (Fig. S6E, S6F). Thus, in contrast to neonatal treatment, T₃ supplementation initiated during adolescence does not rescue social novelty or repetitive behaviors, indicating that normalization of TH signaling must occur within a critical early-life window to reverse *Asxl3*-dependent circuit and behavioral abnormalities.

### Intein-mediated AAV expression of full-length ASXL3 rescues neurodevelopmental deficits in *Asxl3*^+/-^ mice

Because the large coding sequence of *Asxl3* exceeds the packaging capacity of standard AAV vectors, we adopted an intein-based protein-splitting strategy to reconstitute full-length ASXL3 *in vivo*. We used the engineered Cfa_GEP_ split intein, which efficiently mediates protein trans-splicing and ligation^31^, to reconstruct ASXL3 from two separate fragments (Fig. 6A). The *Asxl3* coding sequence was divided into N- and C-terminal segments, each fused to the appropriate intein half and epitope tags, and the two AAV-PHP.eB vectors were co-delivered into neonatal mouse brains via retro-orbital injection (Fig. 6A, 6B).

**Figure 6.**
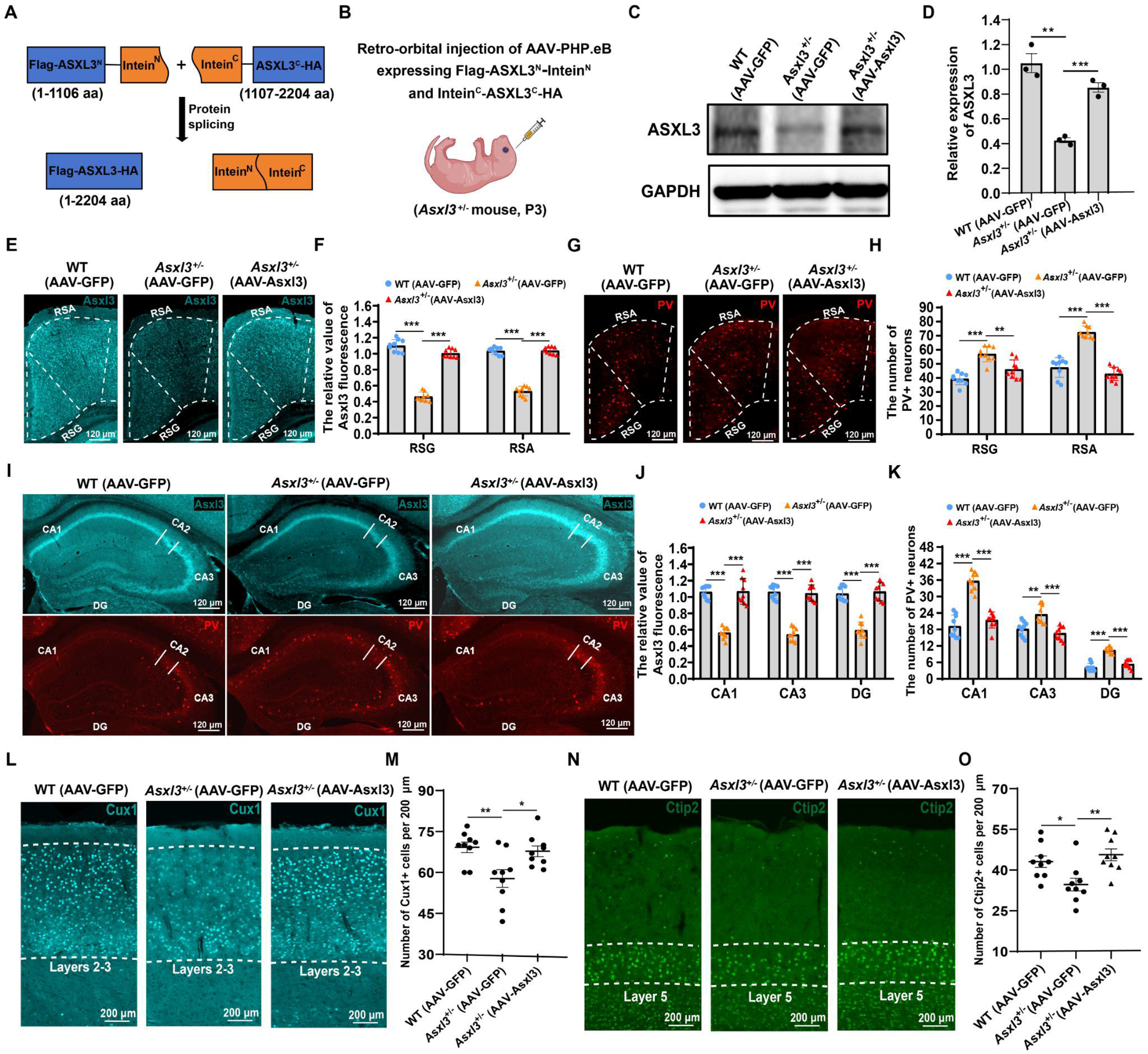
Restoration of *Asxl3* expression reversed cortical and hippocampal neurodevelopmental abnormalities in *Asxl3*^+/-^ mice. (A) Illustration of the molecular strategy for the expression of the intact ASXL3 protein facilitated by the Cfa_GEP_ split intein. (B) Diagrammatic depiction of the retro-orbital injection technique applied to neonatal mice at postnatal day 3. (C) and (D) Immunodetection and quantitative analysis of total ASXL3 protein expression levels. AAV-GFP, injection of AAV-PHP.eB encoding GFP. AAV-Asxl3, injection of AAV-PHP.eB encoding Flag-ASXL3^N^-Intein^N^ and Intein^C^-ASXL3^C^-HA. (E) - (H) Representative images and quantification of ASXL3 and PV expression in the RSC following AAV delivery. (I) - (K) Examination and quantification of ASXL3 and PV expression in the hippocampus after AAV administration. (L) and (M) Immunofluorescence staining of neurons in the second and third cortical layers, accompanied by quantitative analysis of neuronal counts. (N) and (O) Immunofluorescence staining of the fifth layer of cortical neurons and its number statistics. Data are presented as mean ± SD, n = 9 slices from 3 mice for each group. Statistical analysis was performed using one-way ANOVA, with significance denoted by \**P* < 0.05, \*\**P* < 0.01, \*\*\**P* < 0.001.

One month after injection, AAV transduction was widespread throughout the brain, including cortex and hippocampus (Fig. S6G). Western blot analysis using a Flag antibody confirmed robust expression of full-length ASXL3, with an estimated intein splicing efficiency of ∼70% (Fig. S6H, S6I). Total ASXL3 protein levels were significantly increased in the brains of *Asxl3*^+/-^ mice receiving the split-ASXL3 AAVs compared with AAV-GFP controls (Fig. 6C, 6D), indicating successful intein-mediated restoration of ASXL3 expression *in vivo*.

We next asked whether reinstating ASXL3 expression could correct the neurodevelopmental abnormalities observed in *Asxl3*^+/-^ mice. Immunostaining revealed that intein-mediated ASXL3 re-expression normalized PV-positive interneuron number in the RSC (Fig. 6E–6H). In the hippocampus, both ASXL3 and PV expression levels were restored to WT-like values (Fig. 6I–6K), consistent with rescue of inhibitory circuit defects. Moreover, cortical projection neuron deficits were corrected: the reduced numbers of Cux1-positive layer 2/3 neurons and Ctip2-positive layer 5 neurons in *Asxl3*^+/-^ mice were normalized following AAV-Asxl3 treatment (Fig. 6L–6O). Thus, intein-based AAV delivery of full-length ASXL3 is sufficient to reverse the major cortical and hippocampal neurodevelopmental abnormalities caused by *Asxl3* haploinsufficiency.

### Restoration of *Asxl3* expression in *Asxl3*^+/-^ brains rescues ASD-like behavioral phenotypes

We next asked whether intein-mediated restoration of ASXL3, which normalizes cortical and hippocampal development, also ameliorates ASD-like behaviors in *Asxl3*^+/-^ mice. Social behavior was first evaluated using the three-chamber test. *Asxl3*^+/-^ mice injected with AAV-Asxl3 displayed a normalized social novelty response, spending significantly more time interacting with the novel stranger mouse than with the familiar one (Fig. 7A, 7B), in contrast to untreated *Asxl3*^+/-^ mice. Repetitive behavior was assessed using the marble burying test, in which AAV-Asxl3–treated *Asxl3*^+/-^ mice buried fewer marbles than controls (Fig. 7C, 7D), indicating a reduction in repetitive, stereotyped behavior.

**Figure 7.**
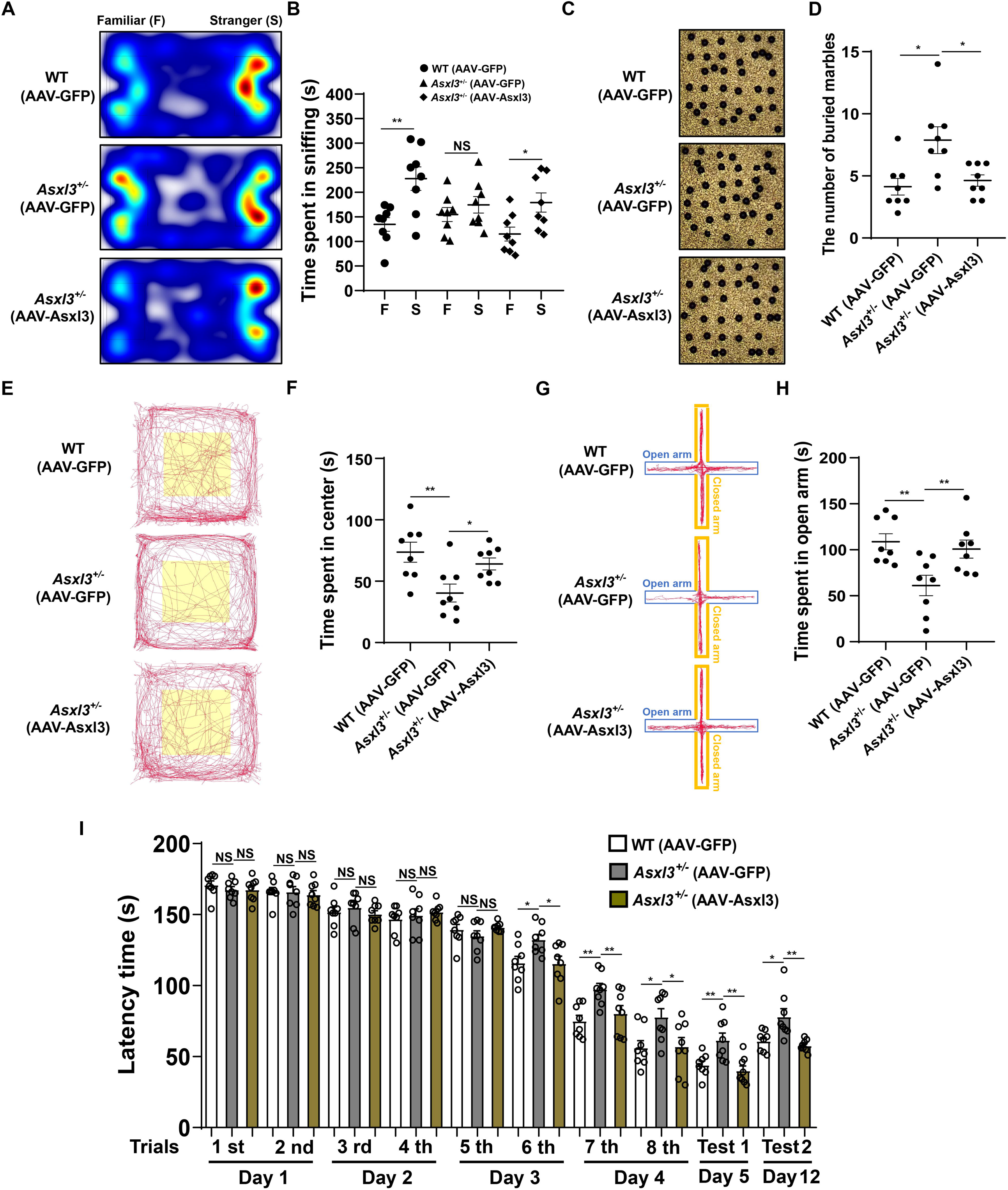
Increasing ASXL3 expression mitigated the ASD-like behaviors in *Asxl3*^+/-^ mice. (A) and (B) Depiction of representative tracing heatmaps and quantification of social interaction time during the social novelty test. (C) and (D) Representative images and statistics of the number of buried beads in the marble burying test. (E) and (F) Representative traces and time statistics for exploring central region in the open field test. (G) and (H) Representative traces and time statistics for exploring open arms in the elevated plus maze test. (I) Statistics on the time it took mice to find the escape box in the Barnes maze test. Data are presented as mean ± SD, n = 8 mice per group. Statistical significance was determined using a two-tailed student’s t-test for Fig. 7B and one-way ANOVA for Figs. 7D, 7F, 7H, 7I. \**P <* 0.05, \*\**P <* 0.01, NS (not significant).

Anxiety-like behavior was then examined. In the open field, AAV-Asxl3–treated *Asxl3*^+/-^ mice spent more time in the central area (Fig. 7E, 7F), and in the elevated plus maze they spent more time in the open arms (Fig. 7G, 7H), consistent with reduced anxiety. Finally, spatial learning and memory were evaluated in the Barnes maze. Starting from the third training day, *Asxl3*^+/-^ mice receiving AAV-Asxl3 showed shorter latencies to locate the escape box compared with *Asxl3*^+/-^ controls (Fig. 7I), indicating improved spatial learning and memory. Together with the anatomical rescue, these data demonstrate that restoring ASXL3 expression via intein-mediated AAV delivery is sufficient to reverse the core ASD-like behavioral abnormalities induced by *Asxl3* haploinsufficiency.

### Intein-mediated expression of full-length ASXL3 is efficient in non-human primate brain

To assess the translational potential of the intein-based ASXL3 restoration strategy, we tested the same split-ASXL3 AAV system in cynomolgus monkeys. AAV9 vectors encoding Flag-ASXL3^N^-Intein^N^ and Intein^C^-ASXL3^C^-HA were administered intrathecally at low or high doses (Fig. 8A). Immunofluorescence staining revealed no detectable Flag signal in the negative control (NC) group, whereas both low-dose and high-dose groups showed robust Flag immunoreactivity in the frontal lobe and hippocampus (Fig. 8B). Higher magnification images confirmed stronger Flag expression in the high-dose compared with the low-dose group (Fig. 8C). Quantitative analysis showed that 80–90% of NeuN-positive neurons in these regions were Flag-positive (Fig. 8D, 8E), indicating widespread neuronal expression of recombinant ASXL3.

**Figure 8.**
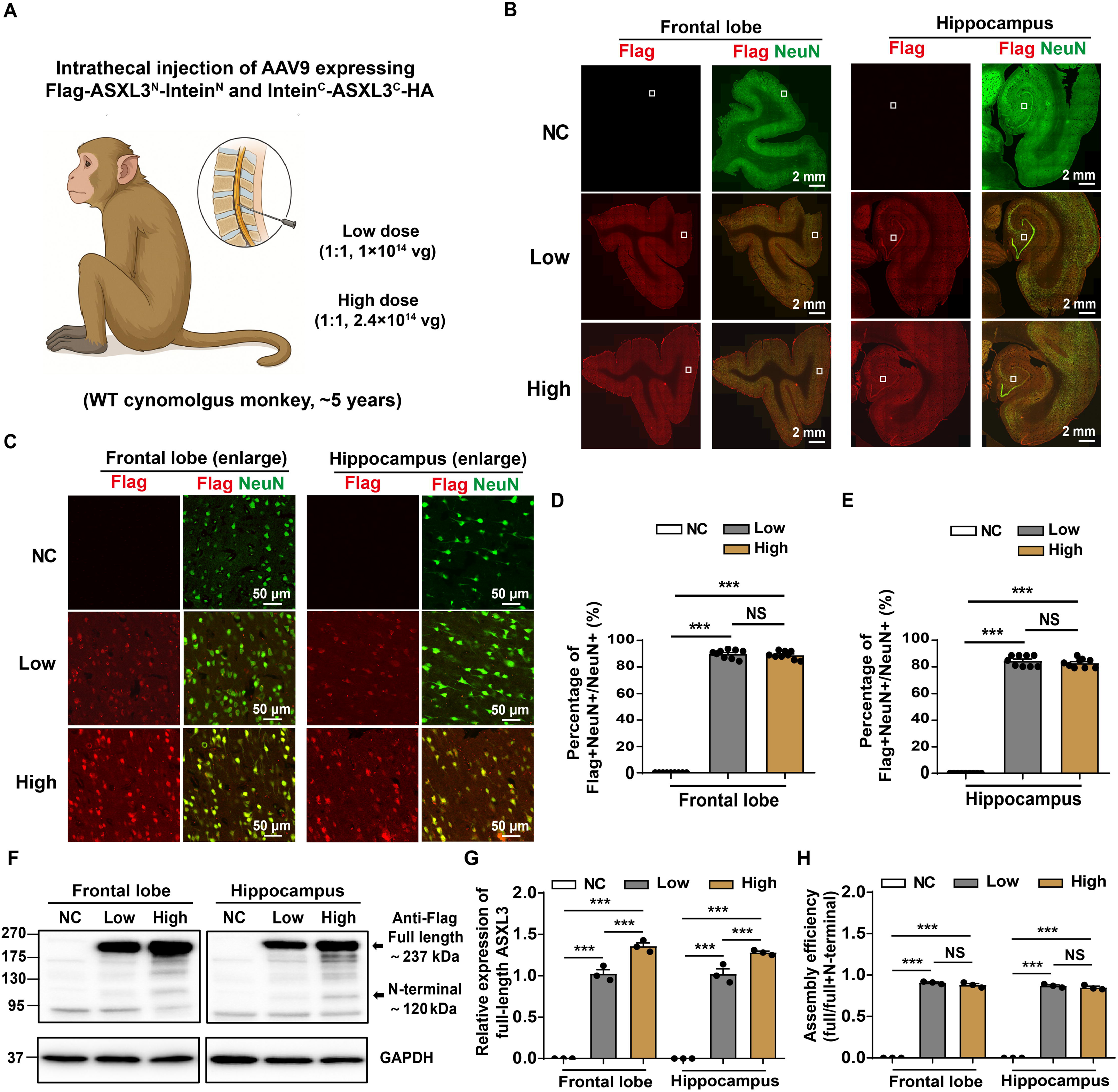
Intein-mediated full-length ASXL3 expression in non-human primates. (A) Schematic diagram illustrating the intrathecal injection-mediated delivery of mixed AAV vectors at two doses (high and low) into the brain of cynomolgus monkeys (B) Representative immunofluorescence images showing Flag and NeuN distribution in the frontal lobe and hippocampus. (C) High-magnification immunofluorescence images of the frontal lobe and hippocampus showing colocalization of Flag with NeuN. (D) and (E) Quantitative analysis of Flag-NeuN colocalization efficiency in the frontal lobe and hippocampus (n = 9 slices from 1 monkey for each group). (F) Detection of recombinant ASXL3 protein expression in the frontal lobe and hippocampus via Western blot. (G) Quantitative analysis of recombinant ASXL3 protein expression levels among different groups (n = 1 monkey per group, with three technical replicates included). (H) Quantitative analysis of intein splicing efficiency (n = 1 monkey per group, with three technical replicates included). The splicing efficiency of the intein was calculated as the ratio of the expression level of full-length ASXL3 protein to the total expression level of full-length ASXL3 and the N-terminal fragment. Data are presented as mean ± SD. Statistical significance was determined by one-way ANOVA. \*\*\**P <* 0.001, NS (not significant).

Western blotting further confirmed efficient reconstitution of full-length ASXL3. In frontal lobe and hippocampal lysates from AAV-treated monkeys, an anti-Flag antibody detected a ∼237 kDa band corresponding to full-length ASXL3 and a ∼120 kDa band representing the N-terminal ASXL3 fragment prior to splicing, and neither band was present in NC samples (Fig. 8F). Quantification revealed that full-length ASXL3 levels were significantly higher in the high-dose group, reaching approximately 1.3-fold those in the low-dose group (Fig. 8G). In both regions, intein splicing efficiency—calculated as full-length ASXL3 divided by total ASXL3 (full-length plus N-terminal fragment)—was ∼90% in both dose groups (Fig. 8H), demonstrating that the intein system maintains high and stable protein-splicing performance in the non-human primate brain across the tested dose range.

## Discussion

ASD is a highly heterogeneous neurodevelopmental condition, and defining pathogenic mechanisms that connect specific gene mutations to circuit- and behavior-level phenotypes remains a major challenge. Here, we show that *Asxl3* haploinsufficiency induces ASD-like behaviors in mice through a previously unrecognized epigenetic–hormonal–circuit axis involving *Dio3* upregulation, reduced TH levels, and abnormal development of PV-positive interneurons. Neonatal TH supplementation or intein-mediated restoration of ASXL3 expression reverses both neurodevelopmental and behavioral abnormalities, highlighting the ASXL3–DIO3–TH–PV interneuron axis as a targetable pathway and providing mechanistic insight into ASD pathogenesis.

As an epigenetic regulator, ASXL3 modulates *Dio3* expression by altering H2A monoubiquitination at the *Dio3* promoter, directly linking chromatin regulation to TH metabolism and PV-positive interneuron development. DIO3 is a TH-inactivating enzyme that is highly expressed in the central nervous system and is essential for maintaining appropriate local TH concentrations during neurodevelopment^32-34^. Our data indicate that *Asxl3* loss increases *Dio3* expression, lowers T₄ and T₃ levels in serum and brain, and disrupts PV-positive interneuron development. Mechanistically, conditional deletion of the TH receptor *Thra* in Nkx2.1-lineage inhibitory neuron progenitors increases PV-positive interneuron density, phenocopying the *Asxl3*^⁺/⁻^ phenotype and establishing TH signaling as a key mediator of the observed interneuron abnormalities. This is consistent with broader evidence that T₃ regulates multiple target genes involved in PV interneuron and interneuron progenitor development^35^. For example, *Shh* loss reduces cortical PV-positive interneuron numbers^36^, whereas *Bmp4* overexpression increases PV-positive cells in the hippocampus^37^. In addition, TH influences *Wnt7a*, which regulates interneuron progenitor proliferation, and *Lhx6*, which is required for interneuron migration and subtype specification^35, 38, 39^.These pathways may contribute to the altered PV-positive interneuron density and cortical layer deficits in *Asxl3*^+/-^ mice. Collectively, our findings define an “ASXL3 → DIO3 → TH → PV-positive interneurons” cascade that bridges genetic mutation, hormonal metabolism, and neural circuit development, providing a concrete example of genetic–metabolic interplay in ASD.

PV-positive interneurons, a major class of GABAergic interneurons critical for maintaining excitation–inhibition balance and network oscillations^40^, have emerged as a convergent cellular substrate in ASD that extends beyond *ASXL3* deficiency. Alterations in PV-positive interneuron number, distribution, or function have been reported across multiple ASD mouse models and in postmortem brains from individuals with ASD^20, 22, 41-46^, supporting their role as a shared pathological node across diverse genetic etiologies. Direct manipulation of PV-positive interneurons or ASD-related genes within these cells has repeatedly demonstrated causal effects on social behavior, repetitive behaviors, and cognition^46-51^, reinforcing the idea that PV-positive interneurons represent a common downstream target disrupted by distinct ASD risk genes. Our work adds a hormonal layer to this framework, showing that epigenetically driven changes in TH metabolism can converge on PV interneuron development. This convergence raises the possibility that interventions aimed at normalizing PV-positive interneuron maturation—such as TH-based strategies—could have broader relevance across ASD subtypes characterized by TH signaling defects or PV interneuron dysregulation.

The therapeutic implications of our findings are twofold. First, neonatal T₃ supplementation in *Asxl3*^+/-^ mice normalizes PV-positive interneuron density and rescues social, repetitive, anxiety-like, and cognitive phenotypes, whereas the same intervention initiated in adolescence is ineffective. This stark contrast underscores a critical early-life window during which TH signaling must be corrected to restore normal circuit development and behavior. Clinically, early TH supplementation could represent a precision therapy for ASD subtypes associated with *ASXL3* mutations or TH metabolic abnormalities. Identifying reliable biomarkers—such as peripheral TH levels, DIO3 expression, or TH-responsive transcriptional signatures—will be essential for stratifying patients most likely to benefit, and careful work will be required to define safe dosing, timing, and potential combination with other interventions.

Second, our intein-based AAV strategy demonstrates that full-length ASXL3 can be efficiently reconstituted *in vivo* despite its large coding sequence. In *Asxl3*^+/-^ mice, intein-mediated ASXL3 restoration corrects PV-positive interneuron density, cortical layering defects, and ASD-like behaviors, establishing ASXL3 replacement as a causal and durable rescue approach. Importantly, the same split-ASXL3 system drives widespread, neuron-enriched expression of full-length ASXL3 with high splicing efficiency in the frontal cortex and hippocampus of non-human primates, supporting the translational feasibility of this gene therapy strategy. Although substantial optimization and safety evaluation will be required before clinical application, these data provide an important proof of concept that large ASD risk genes such as *ASXL3* can be targeted with next-generation AAV–intein platforms.

### Limitations of the study

This study has several limitations. First, our *Asxl3*^+/-^ mouse model relies on partial gene deletion rather than patient-specific single-nucleotide variants, and thus may not fully capture the spectrum of functional consequences and potential gain-of-function or dominant-negative effects associated with human *ASXL3* mutations. Second, species differences in brain development, TH physiology, and PV interneuron circuitry limit direct extrapolation of our findings to humans, and we have not yet validated TH alterations or PV-positive interneuron abnormalities in individuals with *ASXL3* mutations. Third, although our data implicate an ASXL3–DIO3–TH–PV interneuron axis, we did not dissect which downstream TH-responsive pathways (e.g., Shh, Bmp4, Wnt7a, Lhx6) are necessary or sufficient for the observed phenotypes, nor did we explore potential TH-independent roles of *ASXL3* in other neural or non-neural cell types. Finally, while the intein-based AAV strategy showed efficient reconstitution of full-length ASXL3 in mouse and non-human primate brains, the primate experiments were limited in scale, lacked functional and behavioral readouts, and did not systematically evaluate long-term safety, immune responses, or off-target effects of either TH supplementation or ASXL3 restoration, all of which will require extensive future investigation before clinical translation.

## Acknowledgments

This work was supported by the following grants: National Natural Science Foundation of China (#82301334 CD, #82430046 ZQ), National Science and Technology Major Project (2025ZD0214700), Project of Medical Technology Research and Transformation supported by Shanghai Municipal Health Commission (2024ZZ1007), China Postdoctoral Science Foundation (#2021TQ0338, #2022M713236), and Shanghai Post-doctoral Excellence Program (#2021395).

## Author contributions

Z.Q. and C.D. designed this study. C.D. was responsible for the behavioral tests, immunohistochemistry, western blotting, and virus injections. Y.Y., Y.H., and Y.Z. participated in the mouse breeding and genotyping. S.W. conducted experiments related to monkeys. This manuscript was written by C.D. and Z.Q., with valuable input from A.D.

## Ethics statement

This study is approved by the Ethics Committee of Xinhua Hospital, Shanghai Jiao Tong University School of Medicine (XHEC-C-2017-062).

## Animals

All mice (C57BL/6J) utilized in this research were housed under stringent conditions within a pathogen-free facility, adhering to a consistent 12-hour light/dark cycle, with illumination from 7:00 AM to 19:00 PM. The ambient temperature was meticulously regulated between 22 to 25 ℃ to ensure a comfortable living environment. Cages were populated with no more than six mice to guarantee optimal feeding and living conditions.

Nkx2.1-CreER (Cat# C001053) and *Thra*^fl/+^ mice (Cat# S-CKO-17774) were purchased from Cyagen company. To generate *Asxl3*^+/-^ mice, we commissioned BIOCYTOGEN company to knock out exons 5 through 14 of the mouse *Asxl3* gene. The WT band was amplified using specific forward primer (5’-TTAGGAAGGCCTG TAAAGATGTTTC-3’) and reverse primer (5’-CCGAGGAAGCCACAGCTCTTTCC A-3’). For the amplification of knockout band, the forward primer (5’-TTAGGAA GGCCTGTAAAGATGTTTC-3’) and a different reverse primer (5’-TCAATCACAAT TTGTGGGCAAAATC-3’) were used.

For all behavioral tests and cellular/molecular experiments, we utilized both male and female mice, including mice aged 2 to 4 months old. To ensure unbiased results, the behavioral tests were conducted on mice that were age- and sex-matched during the light cycle, with the experimenters being blinded to their genotypes. Before the behavioral tests, the mice were acclimated to human handling over three consecutive days. Additionally, they underwent a 30-minute habituation period in the testing room to adapt to the experimental environment. These measures were implemented to minimize stress and ensure the accuracy and reliability of the test results.

### Three-chamber test

The experimental procedure involved a three-chamber apparatus designed to assess social behaviors in mice, the apparatus was composed of three identical chambers. The test was structured into three distinct 10-minute phases. Initially, during the habituation phase, mice were allowed to explore all chambers freely, thereby familiarizing themselves with the experimental setting. Subsequently, the sociability phase was initiated, introducing an unfamiliar conspecific to the test mouse, enabling the assessment of its innate social preferences. This was followed by the social novelty phase, where a second novel mouse was introduced to evaluate the response of test mouse to new social stimuli. Throughout these phases, the duration of proximity to the introduced mice was meticulously recorded and analyzed using the Noldus EthoVision XT 11.5 software.

### Social intruder test

Before the experiment, the test mouse was fed alone for three days. Then, without food or water, it underwent testing in its familiar cage. The testing involved five trials. In the first, a new mouse (stranger 1) was introduced for three minutes, and their interaction time was recorded. This interaction was defined as when the test mouse initiated contact with stranger 1. The first trial was repeated three times with five-minute breaks. For the final trial, stranger 1 was replaced with a different mouse (stranger 2), and the process was repeated. The interactions were captured by a high-definition camera, and the interaction time was manually measured with a stopwatch.

### Novel object recognition test

The testing procedure took place in a box designed with three equally sized chambers, measuring 40 centimeters in width and 60 centimeters in length. The entire process was divided into three stages: habituation, object cognition, and novel object discrimination. During the initial habituation phase, the test mouse was placed in the box, and empty cages were situated in each of the side chambers. This allowed the mouse to freely roam and explore the environment for a duration of 10 minutes. Next, the object cognition phase commenced. In this phase, a green cone block was placed within the left chamber. The test mouse was then given another 10 minutes to freely explore the box. Finally, the novel object discrimination phase took place. In this phase, a pink cone block (designated as the new object) was placed into the empty cage in the right chamber. The test mouse was given 10 minutes to explore both objects and discern the difference between them.

### Marble burying test

The test was conducted within a sanitized enclosure measuring 40 centimeters in width and length, furnished with a 7-centimeter-thick layer of bedding material. Thirty-six glass marbles were evenly distributed across the bedding surface. The mouse was then allowed to freely navigate and interact with the marbles for a duration of 10 minutes. At the end of the exploration period, the number of marbles that were at least 50% covered by the bedding material was tallied.

### Self-grooming recording

The test mouse was placed into a clean cage devoid of food and water, and its activity within the cage was meticulously recorded for a duration of 30 minutes using a high-definition camera from Da Hua. The grooming behaviors observed during this period encompassed wiping faces, scratching heads and ears, as well as grooming the entire body. These behaviors were carefully documented, and the grooming time was manually tallied using a precision stopwatch.

### Open field test

The test mouse was initially placed in a clean box measuring 40 centimeters in width and length for a habituation period of 10 minutes. Following this acclimation phase, the mouse’s activity was captured using a camera for a subsequent 10-minute duration. The duration spent in the central area of the box, a 20 cm x 20 cm region, was subsequently analyzed using the Noldus EthoVision XT 11.5 software.

### Elevated plus maze

In this test, we used an elevated plus maze with two open and two closed arms, each measuring 6 cm wide and 30 cm long. First, the test mouse was placed in the central area of the maze (6 cm x 6 cm) for 5 minutes to acclimate. Then, we recorded its behavior for another 5 minutes using a camera. Finally, we analyzed the time the mouse spent in the open arms using Noldus EthoVision XT 11.5 software.

### Barnes maze

The Barnes maze, a circular platform with a diameter of 2 meters and 40 small holes around its perimeter, was employed in this experiment. A darkened escape box was concealed beneath one of these holes, while a bright light overhead served as a stimulus for the test mouse to locate the escape box. The study comprised a four-day training period followed by testing on the fifth and twelfth days. During training, the mouse was initially placed in the center of the maze and covered with a black box. Upon removing the box, the mouse was allowed to freely explore for three minutes. The time taken by the mouse to find the escape box was recorded, and once found, it was allowed to rest in the box for a minute. If the mouse failed to locate the escape box within the allotted time, it was still permitted to rest in the box. After each training session, the maze was thoroughly cleaned, and the platform was rotated to position a new hole above the escape box. On the fifth and twelfth days of testing, the time taken by the mouse to find the escape box was again recorded.

### Western Blot

To extract proteins from mouse brain tissue, a homogenization process was carried out using a lysis buffer (Beyotime, P0013) that contained a 1% protease inhibitor cocktail (Beyotime, P0015). Subsequently, the protein concentration was determined by utilizing a BCA Protein Assay Kit (Beyotime, P0010S). For electrophoresis, the extracted proteins were loaded onto a series of 6%-15% BeyoGel™ Plus PAGE gels (Beyotime, P0451, P0455, and P0458). After separation, the proteins were transferred onto PVDF membranes (Millipore, IPVH00010) for immunodetection. The membranes were blocked with 5% milk for 2 hours at room temperature to minimize nonspecific binding. Then, the membranes were incubated overnight at 4°C with primary antibodies. The primary antibodies used are as follows: DIO3 (1:1000, Bioss, bs-3902R), Tubulin (1:5000, abcam, ab7291), Flag (1:1000, abcam, ab205606), HA (1:4000, abcam, ab9110), ASXL3 antibody was made in ABclonal company, and ASXL3 (344-498 aa) region was selected as the antigenic peptide. After washing with PBST to remove unbound antibodies, the membranes were incubated with secondary antibodies for 2 hours at room temperature. Finally, the protein bands were visualized using a chemiluminescent substrate (Thermo Fisher Scientific, 32106).

### Real-Time quantitative PCR (RT-qPCR)

Total RNA was extracted using the RNAsimple Total RNA Kit (TIANGEN, DP419). This RNA was then converted to cDNA with the HiScript III 1st Strand cDNA Synthesis Kit (Vazyme, R312-01). SYBR Green Realtime PCR Master Mix (TOYOBO, QPK-201) was employed to quantitate gene expression, housekeeping gene *Gapdh* was selected as an internal standard. The RT-qPCR cycling conditions were set as follows: an initial denaturation at 95°C for 3 minutes, followed by 40 cycles of denaturation at 95°C for 15 seconds and annealing/extension at 60°C for 60 seconds. Primer sequences used in this study are listed in table S1.

### Immunofluorescence assays

The mice were initially perfused with cold phosphate-buffered saline (PBS) to ensure rinsing of the brain tissues. Subsequently, they were perfused with cold 4% paraformaldehyde to fix the cellular structures within the brain. the brains were excised and immersed in a 4% paraformaldehyde solution at 4°C overnight. Dehydration was achieved through successive immersion in 15% and 30% sucrose solutions for 2 days, respectively. The brains were embedded in OCT compound (SAKURA, 4583) and frozen at −20°C. Using a cryostat (Leica, CM1950), 40-micron-thick slices were cut at −25°C. These sections were then blocked with a solution containing 5% bovine serum albumin (BSA) and 0.3% Triton X-100 in PBS for 2 hours at room temperature. After blocking, the brain sections were incubated overnight at 4°C with primary antibodies. The primary antibodies were as follows: PV (1:500, Abcam, ab181086), Cux1 (1:500, Abcam, ab307821), Ctip2 (1:500, Abcam, ab18465), TBR1 (1:500, Abcam, ab183032), NeuN (1:500, Millipore, ABN78), SST (1:250, Santa Cruz, sc-55565), VIP (1:500, Abcam, 63269), Olig2 (1:500, Millipore, AB9610), Iba1 (1:500, Abcam, ab5076). The sections were then washed three times with PBS to remove unbound antibodies. The sections were incubated with fluorescently labeled secondary antibodies for 2 hours at room temperature. After washing, the immunostained sections were carefully mounted on microscope slides using Fluoromount-G (Southern Biotechnology, 0100-01).

### RNA Sequencing

Total RNA was extracted using Trizol reagent (Magen, R4130). RNA quality was evaluated with the Agilent 2200 system, and samples with an RNA Integrity Number (RIN) greater than 7.0 were considered qualified for subsequent experiments. In accordance with the manufacturer’s protocol, the VAHTS Universal V8 RNA-seq Library Prep Kit (Vazyme, NR605) was used to construct cDNA libraries from the RNA samples. The detailed experimental procedure was as follows: 1 μg of total RNA was taken, and mRNAs with poly-A tails were isolated using oligo(dT) magnetic beads. Subsequently, the mRNAs were fragmented into 200-600 bp segments by treatment with divalent cations at 85°C for 6 minutes. The cleaved RNA fragments served as templates for the synthesis of both the first and second strands of complementary DNA (cDNA). The synthesized cDNA fragments underwent end repair, A-tailing, and ligation with indexed adapters. After purification, the cDNA fragments were treated with uracil DNA glycosylase to remove the second-strand cDNA. The purified first-strand cDNAs were amplified by PCR to generate the cDNA libraries. Library quality was verified using the Agilent 2200 system, and qualified libraries were subjected to 150 bp paired-end sequencing on the NovaSeq 6000 platform. The RNA sequencing data have been deposited to the NCBI Sequence Read Archive (SRA). The accession number is PRJNA1356477.

For RNA sequencing data mapping and analysis, raw sequencing reads were first processed to remove adapter sequences and low-quality reads, yielding high-quality clean reads. The Hisat2 software was employed to align the clean reads to the mouse genome. Additionally, HTseq software was used to calculate gene counts, and the FPKM method was applied to estimate gene expression levels. Finally, the DESeq2 software was utilized to identify differentially expressed genes.

### Chromatin immunoprecipitation (ChIP)

This experiment was conducted using the High-sensitivity ChIP kit (abcam, ab185913). The protocol commenced with the cross-linking of 50 mg of hippocampal and cortical tissues using 1% formaldehyde for a duration of 20 minutes. Subsequently, the tissues were homogenized in 0.5 mL of working lysis buffer to facilitate the release of chromatin. After decanting the supernatant, the chromatin pellet was resuspended in 0.3 mL of ChIP buffer. To fragment the chromatin, sonication was employed using a Bioruptor UCD-200 sonicator. This involved 20 cycles, each consisting of 15 seconds of sonication followed by 30 seconds of rest at a medium power setting. 1 μg of H2Aub antibody (Millipore, 05-678) was added to an assay strip well for 90 minutes at room temperature. 40 μL of the sonicated chromatin was added to the well and incubated overnight at 4°C. The following day, the reaction well was thoroughly washed to remove any unbound material, and the crosslinking was reversed by adding DNA release buffer and proteinase K. The purified ChIP DNA was then resuspended in 20 μL of elution buffer, ready for qPCR analysis. The qPCR primers, specifically designed to amplify the regions of interest, were as follows:

ChIP Primer set 1: forward, CTCCAGTCTCGACGTTCCC; reverse, CACTAGGGGTAGCTGTTGCC.

ChIP Primer set 2: forward, TGAGATCGGGCTCCTAGTCC; reverse, AAATCTTGGGTCGAGGGCG.

ChIP Primer set 3: forward, CTCTCTGCTGCTTCACTCGCT; reverse, CTTGCGGATGCACAAGAAA TCTAA.

### AAV injection

The AAV-PHP.eB-hSyn-Flag-Asxl3^N^-Intein^N^, AAV-PHP.eB-hSyn-Intein^C^-Asxl3^C^-3HA, and AAV-PHP.eB-hSyn-EGFP were procured from PackGene Biotech for retro-orbital injection. We injected 20 μL of mixed AAV (1:1, 1×10^13^ vg/mL AAV-PHP.eB-hSyn-Flag-Asxl3^N^-Intein^N^ and 1×10^13^ vg/mL AAV-PHP.eB-hSyn-Intein^C^-Asxl3^C^-3HA) into neonatal mice (P3). As a control, we injected 20 μL of AAV-PHP.eB-hSyn-EGFP (1×10^13^ vg/mL) into neonatal mice (P3).

The AAV9-hSyn-Flag-Asxl3^N^-Intein^N^ (1×10¹⁴ vg/mL) and AAV9-hSyn-Intein^C^-Asxl3^C^-3HA (1×10¹⁴ vg/mL), also sourced from PackGene Biotech, were administered to cynomolgus monkeys via intrathecal injection. For the low-dose group, 0.5 mL of each virus was injected. For the high-dose group, the injection volume was 1.2 mL per virus.

## Statistical Analysis

The GraphPad Prism 8 software was used to data analysis. The values obtained were expressed as the mean ± standard deviation (SD), providing a comprehensive overview of the variability within each experimental group. For comparisons between two groups, the two-tailed Student’s t-test was employed. When comparing data across multiple groups, one-way ANOVA was utilized to assess any significant variations among the different groups. To indicate the statistical significance of the observed differences, the following notation was adopted: \**p* < 0.05 denoted a significant difference, \*\**p* < 0.01 indicated a highly significant difference, and \*\*\**p* < 0.001 represented an extremely significant difference. In cases where no significant difference was observed, NS (not significant) was used to denote the absence of statistical significance.

