## Supplemental information for "An ASXL3–thyroid hormone axis in parvalbumin interneurons controls autism-like behaviors"

**Supplementary figure 1**

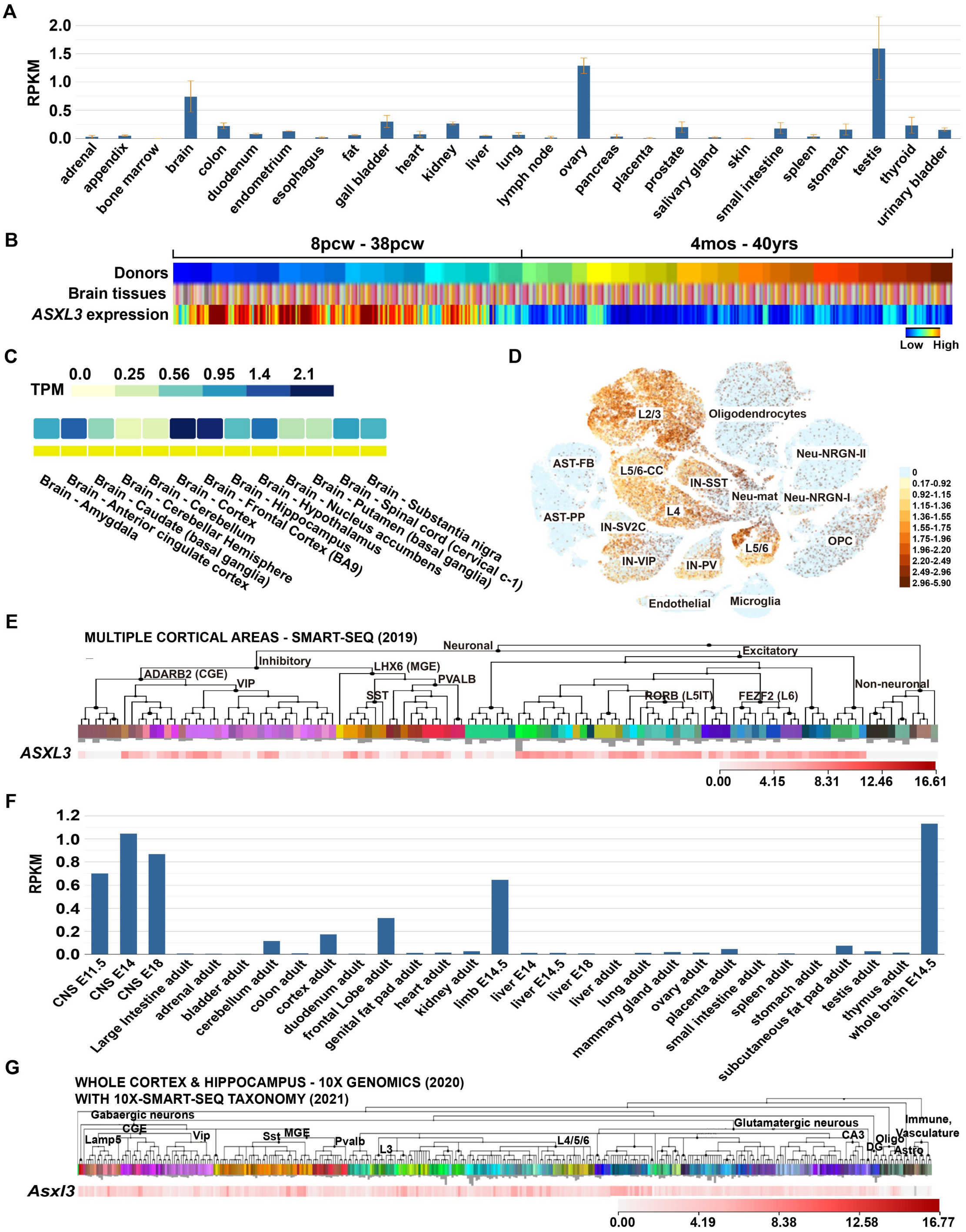


**Figure S1. *ASXL3*/*Asxl3* is enriched in the developing brain and widely distributed across neuronal types**

(A) Comparative expression levels of *ASXL3* in different human tissues. This result is derived from human RNA sequencing data available on the NCBI website (BioProject: PRJEB4337).

(B) Temporal expression profiling of *ASXL3* during human brain development, informed by the BrainSpan database (https:// www. brain span.org/).

(C) *ASXL3* expression across various brain regions, as reported in the GTEx database (https://www.gtexportal.org/).

(D) Single-cell RNA sequencing data from the UCSC cell browser (https://autism. cells.ucsc.edu), providing insights into *ASXL3* expression across diverse neuronal cell types.

(E) Single-cell RNA sequencing data from the Allen Brain Map, further demonstrating *ASXL3* expression in diverse neuronal populations.

(F) *Asxl3* expression across various tissues and developmental phases in mice. This result is derived from mouse RNA sequencing data available on the NCBI website (BioProject: PRJNA66167).

(G) Single-cell RNA sequencing data from cortical and hippocampal tissues, highlighting *Asxl3* expression in diverse neuronal populations. This result is derived from Allen Brain Map.

**Supplementary figure 2**

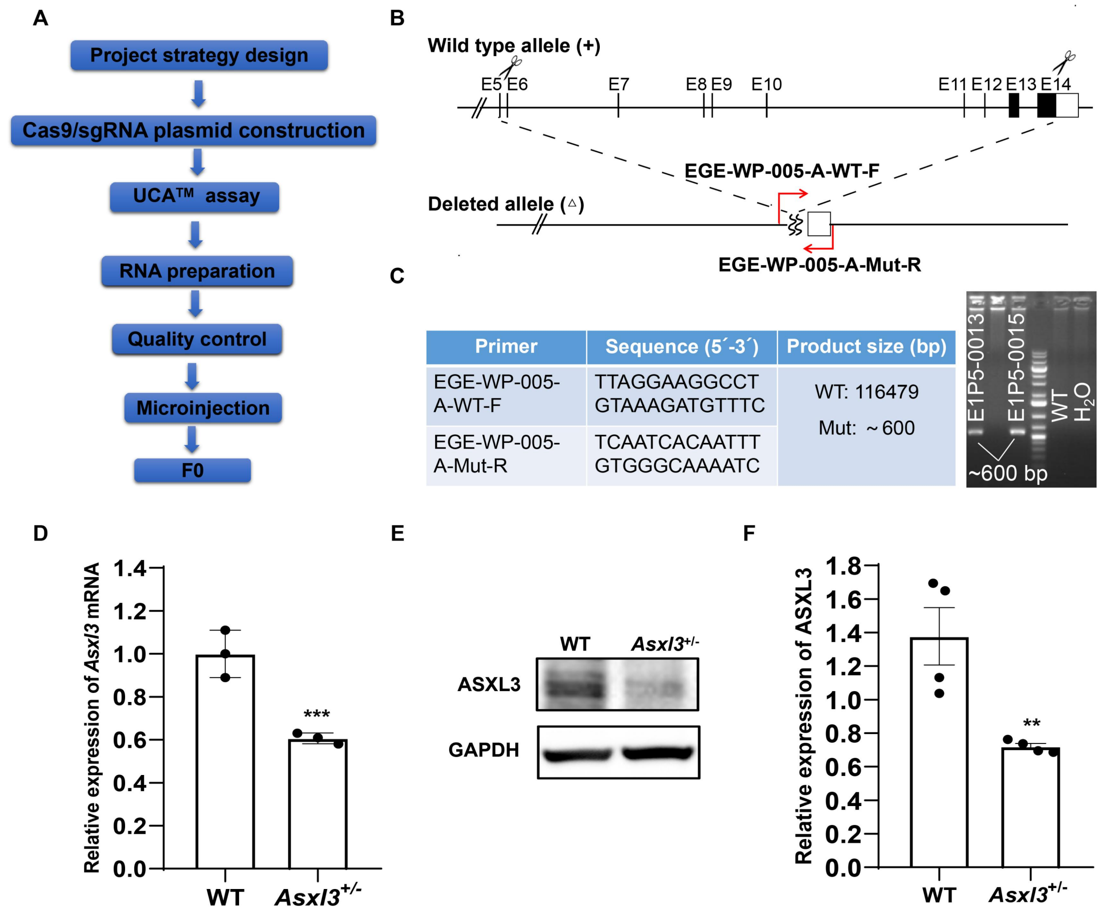


| Primer | Sequence (5´-3´) | Product size (bp) |
|---|---|---|
| EGE-WP-005-A-WT-F | TTAGGAAGGCCT GTAAAGATGTTTC | WT: 116479 Mut: ~600 |
| EGE-WP-005-A-Mut-R | TCAATCACAATTT GTGGGCAAAATC | |

**Figure S2. Generation of *Asxl3* knockout mice via CRISPR/Cas9-mediated genome editing**

(A) Overview of the CRISPR/Cas9-driven strategy for producing heterozygous *Asxl3* mutant mice.

(B) A graphical representation of the targeted deletion of exons 5 to 14 within the *Asxl3*.

(C) Identification of *Asxl3* mutant mice through polymerase chain reaction (PCR).

(D) Quantitative analysis of *Asxl3* mRNA expression in the WT and *Asxl3*$^{+/-}$ mice (n = 3 for both WT and *Asxl3*$^{+/-}$ mice).

(E) Protein-level analysis of *Asxl3* expression in the WT and *Asxl3*$^{+/-}$ mice.

(F) Quantitative analysis of protein expression data (n = 4 for both WT and *Asxl3*$^{+/-}$ mice).

Data are presented as mean ± SD, two-tailed student's t-test, $**P < 0.01$, $***P < 0.001$.

**Supplementary figure 3**

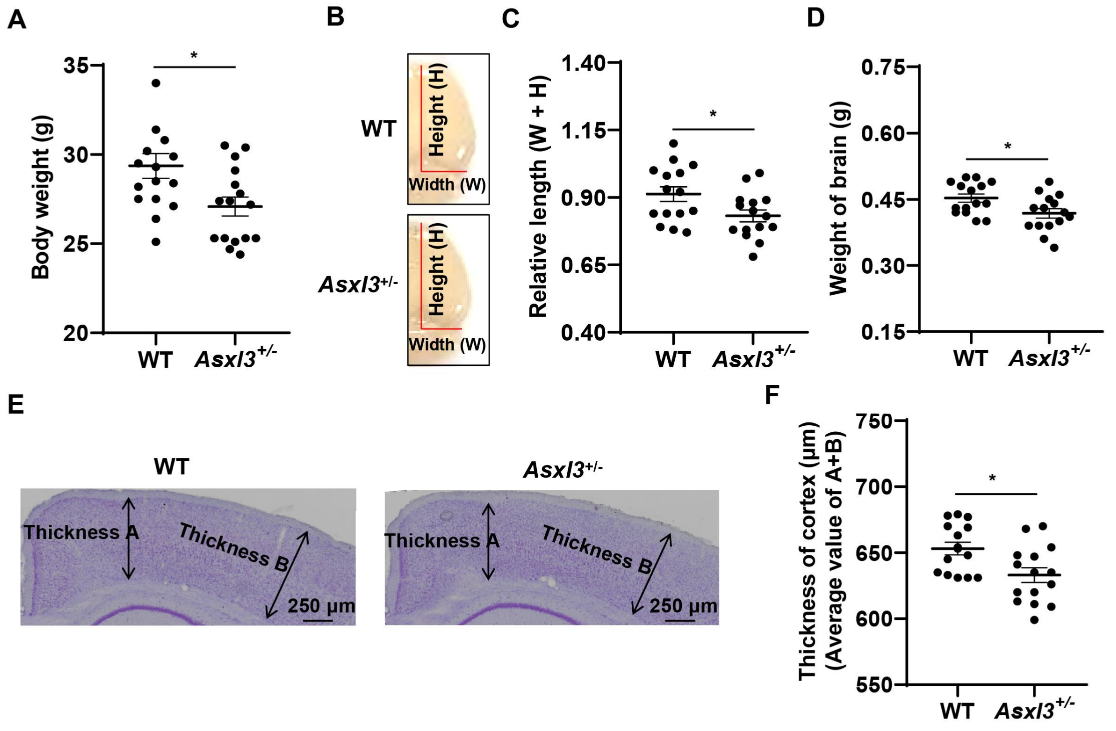


**Figure S3.** ***Asxl3***$^{+/-}$ **mice exhibited abnormal development**

(A) Comparative body weights of 3-month-old WT and *Asxl3*$^{+/-}$ mice (n = 16 per group).

(B) and (C) Graphical representation and statistical analysis of brain dimensions in WT and *Asxl3*$^{+/-}$ mice aged 3 months (n = 15 per group).

(D) Comparative brain weights of 3-month-old WT and *Asxl3*$^{+/-}$ mice (n = 15 per group).

(E) and (F) Nissl staining and cortical thickness measurements in WT and *Asxl3*$^{+/-}$ mice (n = 15 slices from five mice).

Data are depicted as mean ± SD, with statistical significance determined by a two-tailed student's t-test, *$P < 0.05$.

**Supplementary figure 4**

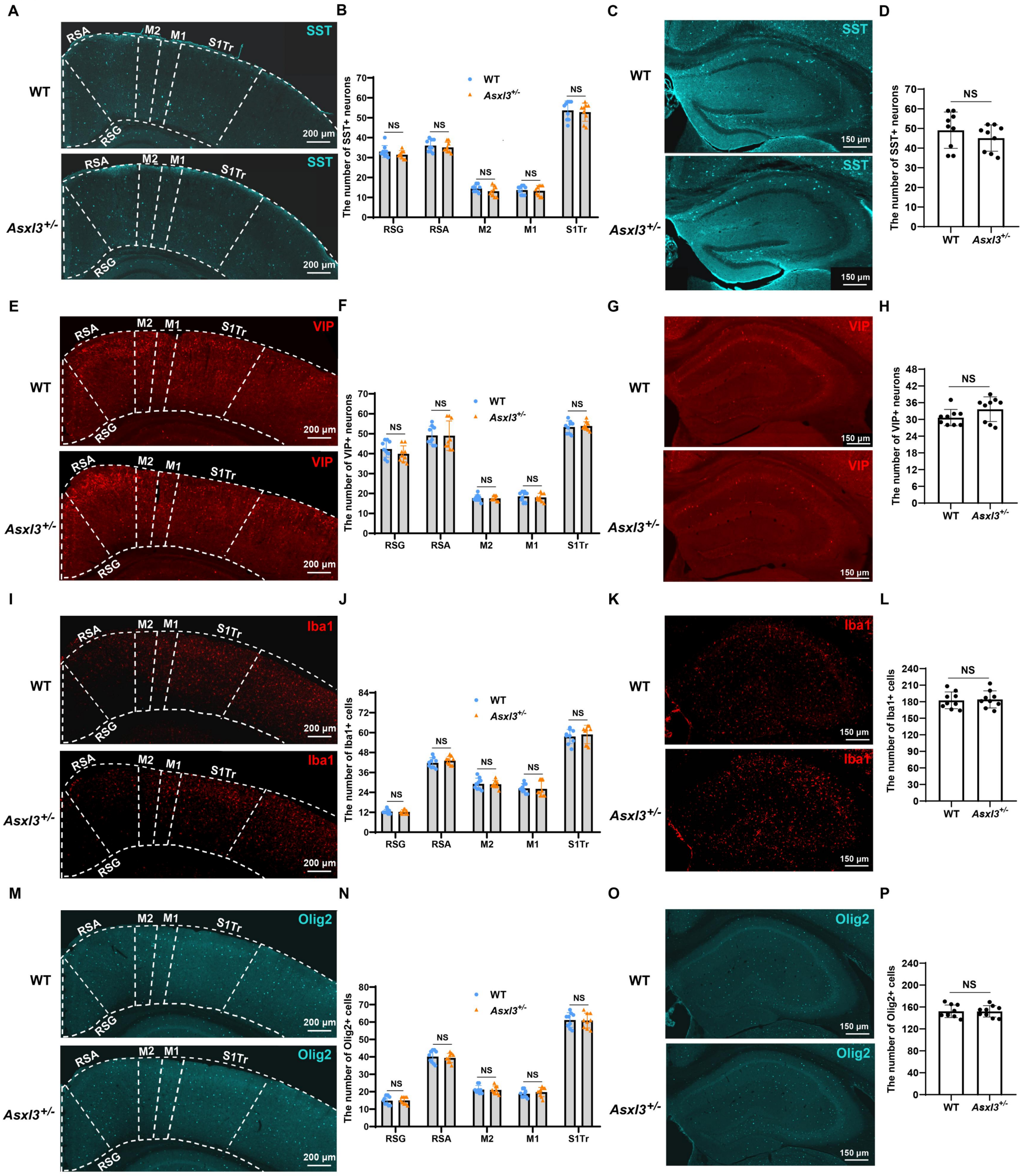


**Figure S4.** ***Asxl3*** **haploinsufficiency did not affect the development of SST-positive neurons, VIP-positive neurons, and glial cells**

(A)-(D) Immunofluorescence staining delineates the distribution and quantification of SST-positive neurons within the cortical and hippocampal regions.

(E)-(H) Immunostaining and number statistics of VIP-positive neurons in the cortex and hippocampus.

(I)-(L) Immunofluorescence staining employing Iba1 as a microglial marker, coupled with a quantitative evaluation of microglial presence in the cortex and hippocampus.

(M)-(P) Immunofluorescence staining with Olig2 as an oligodendrocytic marker, followed by a tally of oligodendrocyte counts in the cortical and hippocampal areas. Data are presented as mean ± SD, n = 9 slices derived from 3 mice per group, statistical analysis conducted using a two-tailed student's t-test, NS (not significant).

Supplementary figure 5

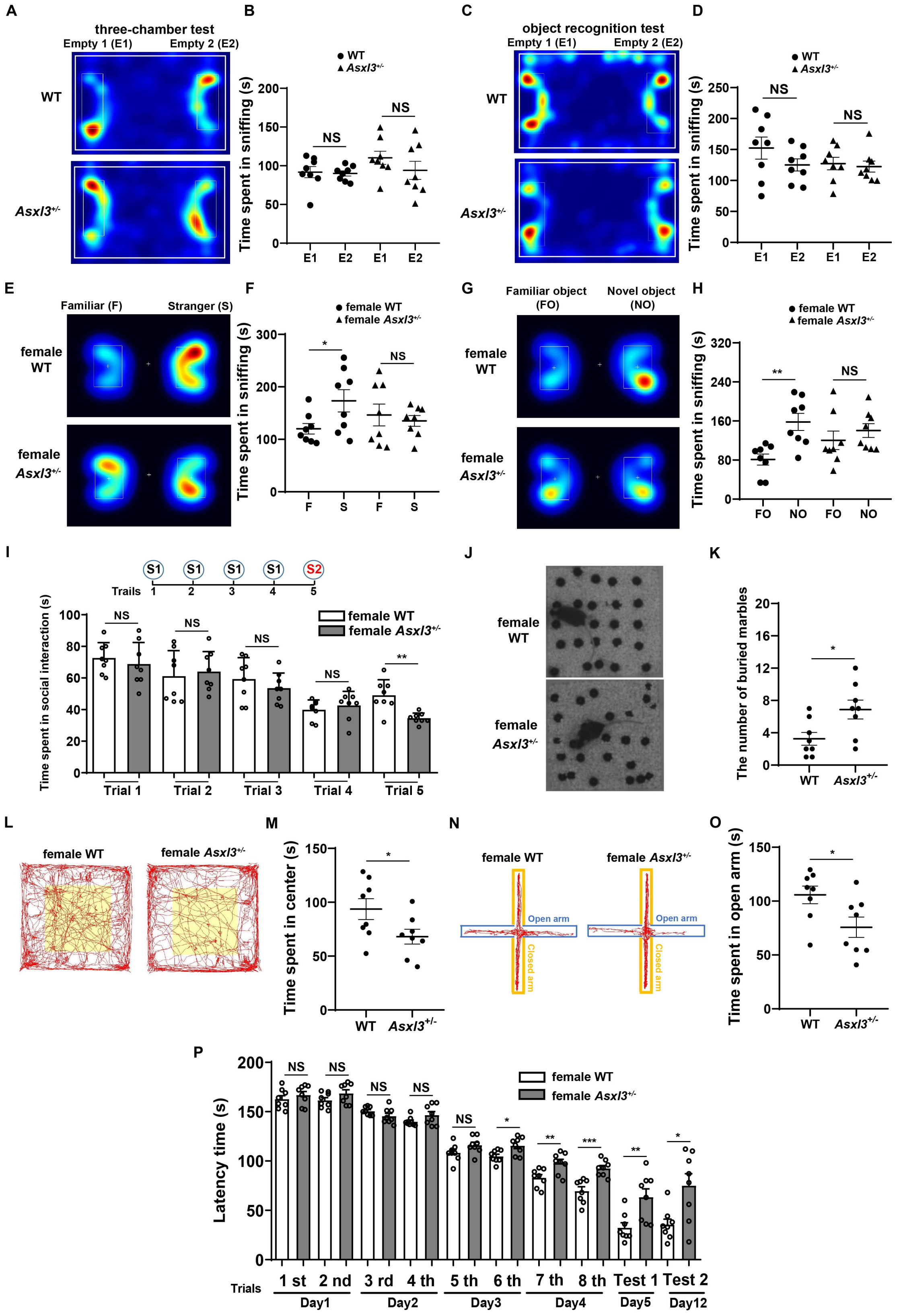


### Figure S5. The female *Asxl3*$^{+/-}$ mice displayed ASD-like behaviors

(A) and (B) Representative heatmaps and quantification of exploration time during the habituation phase of the three-chamber test.

(C) and (D) Representative heatmaps and quantification of exploration time during the habituation phase of the novel object recognition test.

(E) and (F) Representative heatmaps and quantification of social interaction time in the social novelty test.

(G) and (H) Representative heatmaps and quantification of exploration time in the novel object recognition test.

(I) Quantification of social interaction time in the social intruder test.

(J) and (K) Representative images and statistics of the number of buried marbles in the marble burying test.

(L) and (M) Representative images and quantification of time spent exploring the central area in the open field test.

(N) and (O) Representative images and statistics of time spent in open arms of the elevated plus maze test.

(P) Statistics on the time taken by mice to find escape holes in the Barnes maze test.

All data are presented as mean ± SD, n = 8 (WT) and n = 8 (*Asxl3*$^{+/-}$), two-tailed student's t-test, *$P < 0.05$, **$P < 0.01$, ***$P < 0.001$, NS (not significant).

#### Supplementary figure 6

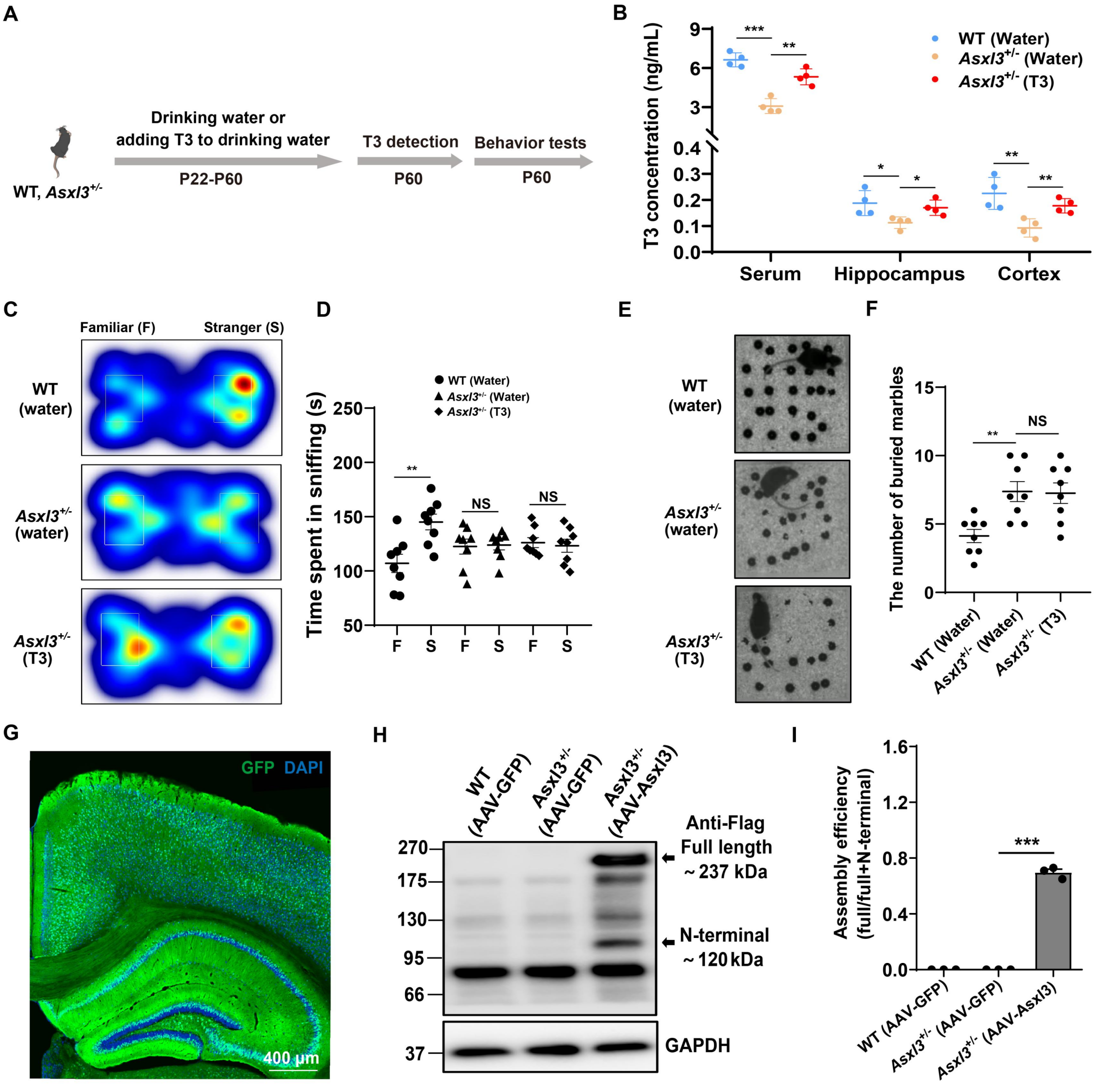


**Figure S6. Supplementation of T3 in adolescent *Asxl3*$^{+/-}$ mice failed to ameliorate ASD-like behaviors, and the efficiency of intein splicing was detected**

(A) The flowchart of T3 supplementation and behavioral tests in adolescent *Asxl3*$^{+/-}$ mice. The concentration of the T3 solution is 0.5 μg/mL.

(B) The expression levels of T3 in the serum and brain of *Asxl3*$^{+/-}$ mice after T3 supplementation, as determined by ELISA (n = 4 mice per group).

(C) and (D) Heat maps of social novelty test results and statistical analysis of social interaction time (n = 8 mice per group).

(E) and (F) Images of the marble burying test results and the statistics of the number of marbles buried (n = 8 mice per group).

(G) Detection of GFP signals in the brain of *Asxl3*$^{+/-}$ mice subsequent to AAV injection.

(H) Western blot analysis was performed to detect the expression of full-length ASXL3 protein and N-terminal protein in mice following AAV injection.

(I) Statistical analysis of intein assembly efficiency (n = 3 mice per group).

Data are presented as mean ± SD. Statistical significance was determined using a two-tailed student's t-test for Fig. S6D and one-way ANOVA for Figs. S6B, S6F, S6I. *$P$ < 0.05, **$P$ < 0.01, ***$P$ < 0.001, NS (not significant).

**Table S1. The information of primer sequences in RT-qPCR assay**

| ***Gene*** | **Forward primer (5' to 3')** | **Reverse primer (5' to 3')** |
|---|---|---|
| *Asxl3* | GCGTTGGTGTGTATGAAGGC | TCCCCGACTCGAGTGTTAGT |
| *Capn 11* | GACTTCTCTCCCACGGACAT | AGGTGGTAAGGGAGCCAATG |
| *Gpr101* | AAGGTGGTCAGAGGAAACGC | AAGTCAGACAGGCACGACAG |
| *Banp* | ATTGGGGAGGATGGACAGGT | CAATGTGGAGGTGGCCCTG |
| *Scn4b* | AACCGAGGCAATACTCAGGC | GCAGGATCGATGAGCCGTTA |
| *Tmem255a* | GCAGATGCTGGTGGCTTCTA | ACATAGTGGCACCGGTTAGC |
| *Arrdc2* | CCGGAAGGTGTTCACTGTCA | AGTGTAGCCTTTGCGGTCAA |
| *Dio3* | TCAGACGACAACCGTCTGTG | GAAGCCATCAGGTCGGACAA |
| *Septin4* | GGACCATTCACTGGGATGGC | TGAGTGGTAGGGTTCGCTTC |
| *Hspa1a* | CACCATCGAGGAGGTGGATT | GACAGTCCTCAAGGCCACAT |
| *Unc5d* | ATGGGGGCAAGTTCTGTGAA | CCTCTCCTACTCCATCTTTGGG |
